# Experimental and field evidence indicate that islet-nesting tundra birds experience reduced nest predation and benefit indirectly from high snow goose densities

**DOI:** 10.1101/2025.11.26.690446

**Authors:** Marylou Beaudoin, Andréanne Beardsell, Éliane Duchesne, Madeleine-Zoé Corbeil-Robitaille, Pierre Legagneux, Dominique Berteaux, Joël Bêty

## Abstract

Landscape features can shape the occurrence and strength of predator-prey interactions by influencing predation risk and prey distribution. In the High Arctic, some bird species select nesting sites with physical features that limit the access by their main terrestrial predator, the arctic fox, though these features do not always provide full protection. We investigated how nest microhabitat characteristics and prey availability modulate nest survival in tundra birds that select pond and lake islets as breeding sites. Over four summers, we analyzed the survival of 132 cackling goose and 55 glaucous gull nests located on islets or on pond and lake shores within a 150 km^2^ area occupied by a snow goose colony on Bylot Island, Nunavut, Canada. We also analyzed survival of 537 artificial nests deployed over three summers. We found that islets act as partial prey refuges, with higher nest survival rates on islets than on pond and lake shores. Nest survival generally increased with islet distance from shore, but we found little evidence of this effect for cackling geese and glaucous gulls, which avoided nesting on islets near shore. Moreover, water depth surrounding islets had little to no influence for any nest type. Nest mortality was much higher in a year with relatively low snow goose nest density, suggesting a short-term positive indirect effect of this colonial nesting bird on species nesting on islets. Since the arctic fox was virtually the sole predator of artificial nests, our findings indicate that annual variation in nest survival on islets were driven by a shift in fox foraging behavior in response to changes in prey availability across the landscape. Our study, which integrates multi-year monitoring and field experiments, highlights the interplay between microhabitat selection and predator-multi-prey dynamics in the arctic tundra.

## Introduction

Physical characteristics of the landscape can shape species relative abundance by influencing the occurrence and strength of interactions (Cherif et al. 2024). Landscape features can affect predator foraging behavior by modulating movement capacity, prey encounter, prey detection, attack probability, and success rate (Shepard et al. 2013; Wootton et al. 2023; Beardsell et al. 2025). These factors collectively influence predation risk for prey (Atuo and O’Connell 2017), ultimately impacting prey distribution and persistence in the landscape (Lima 1998; Laundré, Hernández, and Ripple 2010; Clermont et al. 2021).

Prey can select microhabitats that minimize the likelihood of being predated (Gaynor et al. 2019). These microhabitats, known as prey refuges, can enhance prey survival or reproduction compared to the surrounding habitat (Berryman and Hawkins 2006; Duchesne et al. 2021). The protective effect of refuges can arise from the limited use of these microhabitats by predators due to elevated short- and long-term fitness costs. These costs include reduced predation efficacy and efficiency (e.g., lower attack success, increased cost of transport), and a higher risk of injury (e.g., greater difficulty evading prey defenses, falling from steep terrain) (Mukherjee and Heithaus 2013; Shepard et al. 2013). The quality of refuges can vary, offering complete or partial protection depending on differences in their physical characteristics that influence the foraging behavior of predators and, thus, the associated predation risk (Velando and Márquez 2002; Brown and Kotler 2004). Beyond physical characteristics, the quality of partial refuges may also be context-dependent (Lecomte et al. 2008; Batbayar et al. 2014). For instance, fluctuating local prey densities could modulate the energetic state of predators and, as a result, their willingness to pursue prey located in less accessible or riskier microhabitats (Berger-Tal et al. 2009). Integrating the interplay between physical landscape features and prey densities is therefore crucial for evaluating the quality of refuges and their role in shaping prey distribution and abundance in the landscape (Clermont et al. 2021; Duchesne et al. 2021; Beardsell et al. 2025).

The Arctic tundra provides an excellent system to study how the quality of refuges for prey can be modulated by both the physical characteristics of microhabitats and the food resources available to predators. The arctic fox (*Vulpes lagopus*) is the main nest predator of tundra bird species, heavily influencing their reproductive success (Bêty et al. 2002; McKinnon and Bêty 2009). Its foraging behavior is strongly shaped by annual prey abundance, which can lead to fluctuations in predation pressure on nests (Bêty et al. 2002; Beardsell et al. 2022). Some bird species less effective in protecting their nest against foxes can select islets in ponds and lakes as nesting sites, where water barriers hinder fox access, improving reproductive success compared to shore sites (Zoellick et al. 2004; Gauthier et al. 2015). Although foxes are capable swimmers (Strub 1992), their attack and success probabilities are likely lower for nests on islets because they cannot achieve the same speed or adopt the same offensive or defensive positions as they could on shore (Beardsell et al. 2025). Moreover, foxes must weigh the risk of injury posed by defensive birds, as even seemingly minor injury could prove life-threatening (Mukherjee and Heithaus 2013).

The quality of islets as refuges in the tundra landscape may depend on their physical characteristics. For instance, glaucous gulls (*Larus hyperboreus*), cackling geese (*Branta hutchinsii*), and red-throated loons (*Gavia stellata.*) tend to nest on islets farther from shore and surrounded by deeper water (Corbeil-Robitaille et al. 2024), potentially to reduce predation risk from terrestrial predators (Gauthier et al. 2015). Furthermore, the relative protection offered by islets may also vary depending on the density of prey available to arctic foxes (Gauthier et al. 2015). In most tundra ecosystems, foxes heavily rely on lemmings, whose populations undergo large-amplitude cycles (Giroux et al. 2012; Fauteux, Gauthier, and Berteaux 2015). When lemming abundance is low, predation pressure on nesting birds generally increases (Summers, Underhill, and Syroechkovski 1998; Bêty et al. 2001; Beardsell et al. 2022). Colonial-nesting birds, like snow geese, can also constitute a significant portion of the diet of arctic foxes, primarily through the consumption of their eggs and chicks (Giroux et al., 2012). Fox density is increased in these large bird colonies due to a reduction in their home range size, thereby increasing predation pressure on vulnerable tundra-nesting birds (Dulude-de Broin et al. 2023).

Through field observations and experiments, we assessed the effectiveness of islets as refuges for nesting birds in the tundra ecosystem of Bylot Island (Nunavut, Canada), where lemmings and colonial snow goose populations exhibit strong inter-annual variation in abundance. We examined how nest microhabitat (islet or shore), islet characteristics (distance to shore and water depth) and main prey densities influence nest survival of cackling geese and glaucous gulls. Unlike snow geese, individuals from these species are commonly found on islets, but also actively defend their nests with aggressive behavior, creating a potential risk of injury for foxes (Beardsell et al. 2025). We also conducted field experiments using artificial nests to provide a standardized measure of predation risk that excludes the potential effects of nest defense ability and parental quality (McKinnon et al. 2010). We hypothesized that both natural and artificial nest survival would be higher for nests less accessible to foxes, predicting (P1) greater survival on islets than on shore, and (P2) increased survival on islets with greater distance from shore and water depth, as these factors could increase risks for foxes and reduce their predation efficiency. As foxes are more likely to target prey in less accessible microhabitats when their energy acquisition rate is lower (Beardsell et al. 2025), we also predicted that (P3) nest survival would decrease in years of low lemming and low snow goose nest densities.

## Material and methods

### Study system

We conducted fieldwork in summers 2018, 2019, 2022, 2023 and 2024 in a ∼150km^2^ study area located on the southwest plain of Bylot Island (72°88’N, 79°84’W), in Sirmilik National Park, Nunavut, Canada (Fig. 1). This area is mainly characterized by mesic tundra and polygonal wetlands interspersed with streams, shallow ponds, and lakes (Gauthier et al. 2024). A total of 355 islets have been geolocated in these small waterbodies, with the majority (>80%) remaining unoccupied by birds during the summer (Corbeil-Robitaille et al. 2024). Red-throated loons (*Gavia stellata*), glaucous gulls and cackling geese are the main species selecting islets as nesting sites (Corbeil-Robitaille et al. 2024) (Fig. 2).

**Fig. 1.**
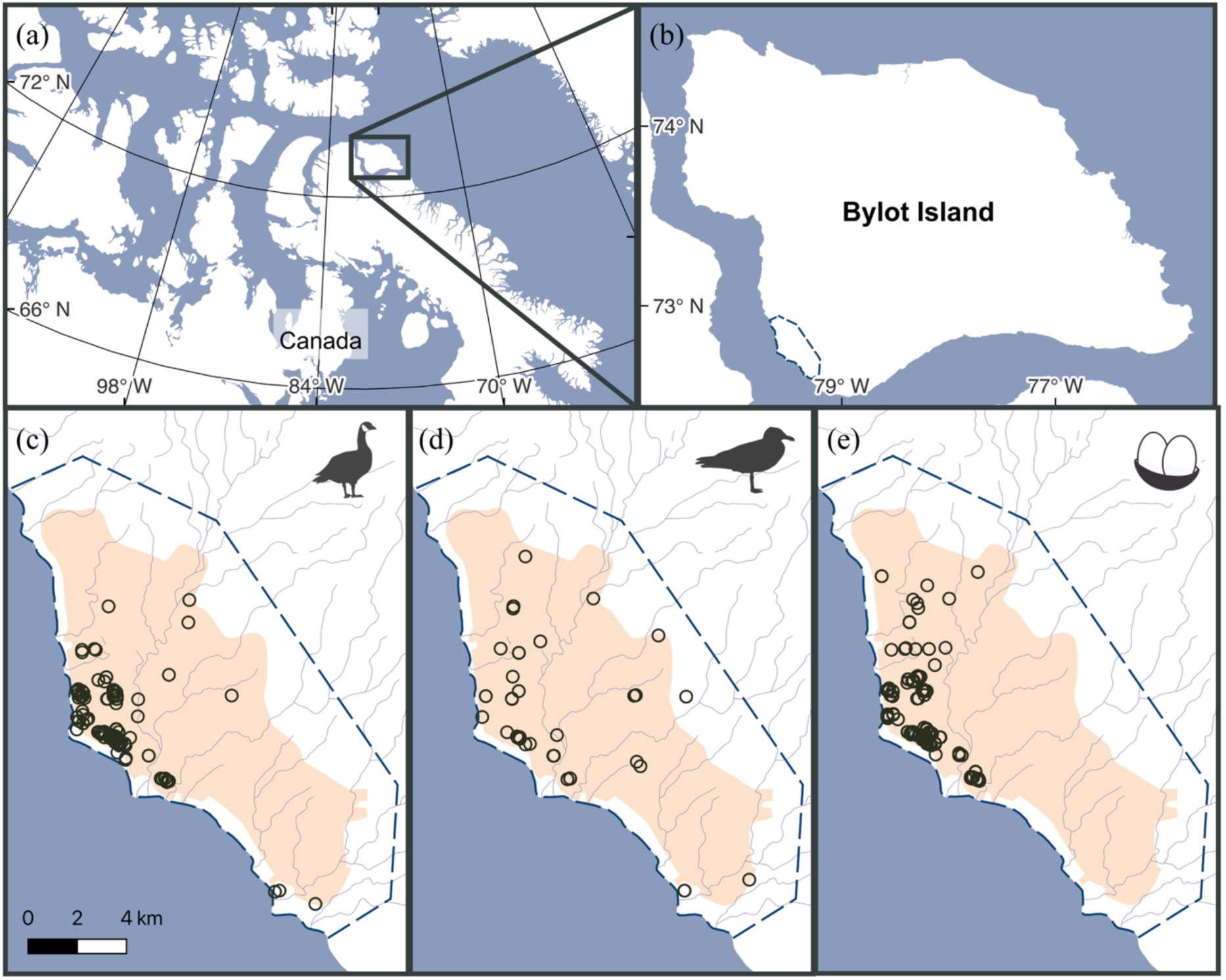
Top panels (a,b) show the study location on Bylot Island, Nunavut, Canada. Bottom panels show the study area (outlined by dashed lines; ∼150 km^2^), including the nesting snow goose colony (orange polygon) and the spatial distribution of nests (hollow circles) used in survival analyses: (c) cackling goose nests (n =132), (d) glaucous gull nests (n = 55) and (e) artificial nest experimental units (n = 179), each consisting of a triad of artificial nests (see Methods).

**Fig. 2.**
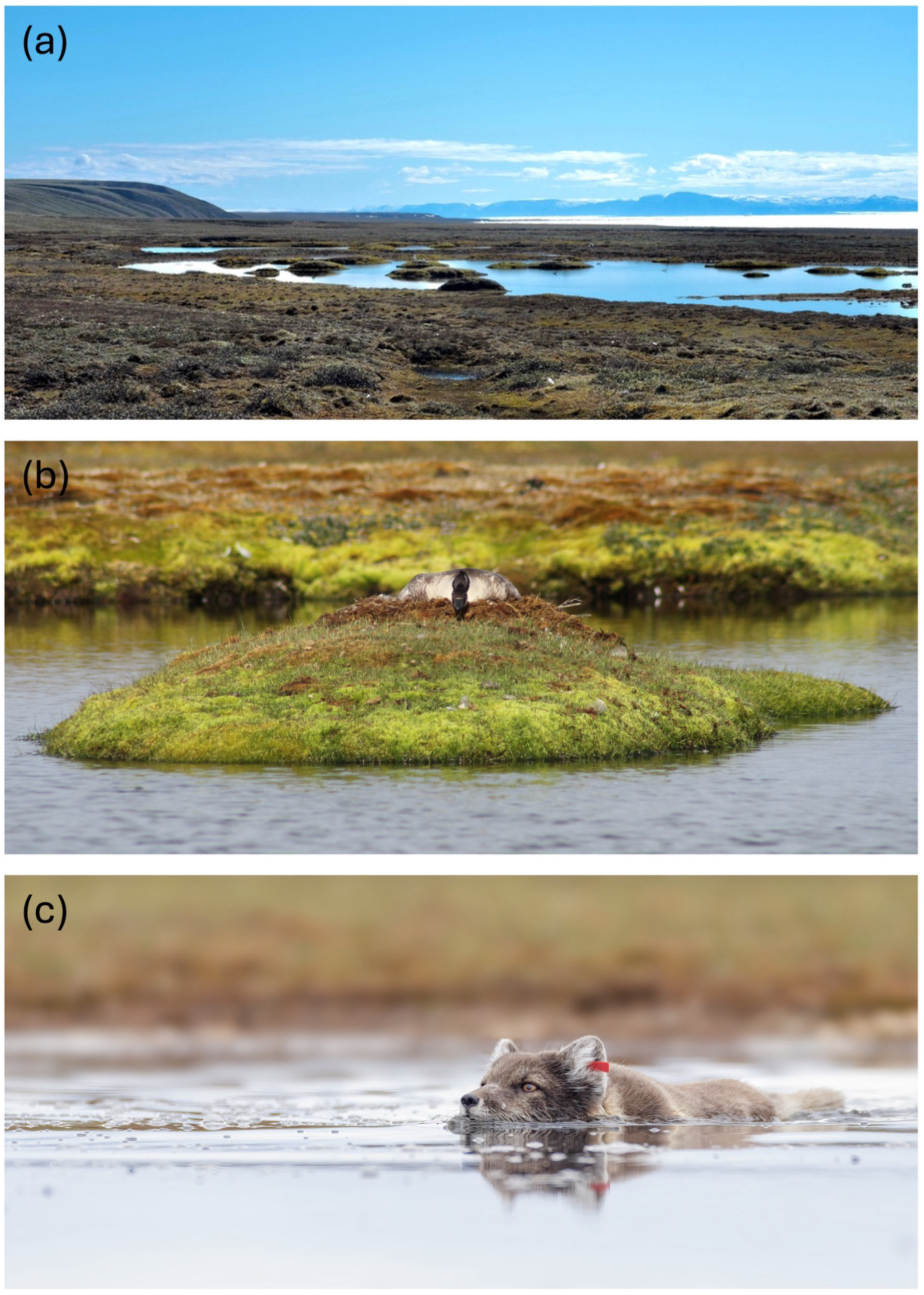
The wetlands of the study area comprise numerous ponds and lakes scattered with islets (a), which serve as nesting sites for a few bird species, including the cackling goose (b). Arctic foxes can reach these islets by swimming (c) or by jumping when the islet is located close to the shore. Photo credits: (a) Jeanne Clermont, (b) Yannick Seyer, and (c) Louis-Pierre Ouellet.

Vertebrate biodiversity is dominated by birds, with over 35 species observed nesting (Moisan et al. 2025). The greater snow goose is the most abundant avian species in the study area, typically breeding each summer in a vast colony of approximately 25,000 pairs across an area of about 75 km^2^ (Moisan et al. 2025) (Fig. 1). The snow goose colony provides an abundant food source for predators, and its average size and spatial extent have remained relatively stable over the past decades (Duchesne et al. 2021; Moisan et al. 2025). However, in 2022, due to a very late spring and delayed snowmelt, the colony reached an unprecedented low, with only ∼4,500 pairs nesting over a reduced area of 40 km^2^ (Moisan et al. 2025). In parallel, predators in the system also heavily rely on small mammals. The brown (*Lemmus trimucronatus*) and collared lemming (*Dicrostonyx groenlandicus*) show synchronized abundance cycles of 3-5 years, with much greater amplitude in the brown lemming’s cycle (Gruyer, Gauthier, and Berteaux 2008).

The arctic fox is the main terrestrial predator in the study area, feeding heavily on lemmings and snow geese (primarily eggs and goslings) during the summer (Bêty et al. 2002; Gauthier et al. 2011). Foxes also prey on the eggs of various nesting birds, including glaucous gulls and cackling geese. Avian predators such as glaucous gulls, long-tailed and parasitic jaegers (*Stercorarius longicaudus* and *S. parasiticus*) and common ravens (*Corvus corax*) may occasionally depredate eggs from undefended nests (Bêty et al. 2001; McKinnon and Bêty 2009). However, their influence on the breeding success of cackling geese and glaucous gulls is likely limited, as these avian predators rarely displace large birds from their nests (Inglis 1977).

### Study design

We analyzed natural and artificial nest survival to assess islet refuge protective quality based on physical characteristics and biological context. Between microhabitats, we compared nest survival on islets and on pond and lake shores for cackling geese (Fig. 3a) and artificial nests (Fig. 3b). We could not conduct this analysis for glaucous gulls as only three nests were monitored on the shore (see Supporting information for annual sample sizes). Between islets, we evaluated how nest survival was influenced by distance from shore and water depth surrounding islets for cackling geese (Fig. 3c), glaucous gulls (Fig. 3d) and artificial nests (Fig. 3e). For natural nests, all analyses included both main prey annual densities (lemmings and snow goose nests). Natural nest monitoring and robust prey density estimates were unavailable for 2024 due to logistical constraints that prevented us from conducting fieldwork during the birds’ incubation period. However, we still deployed artificial nests that year. Consequently, we could not include annual main prey density estimates in artificial nest analyses but nonetheless tested for annual variation in nest survival (Fig. 3b,e), as we could still infer general main prey availability.

**Fig. 3.**
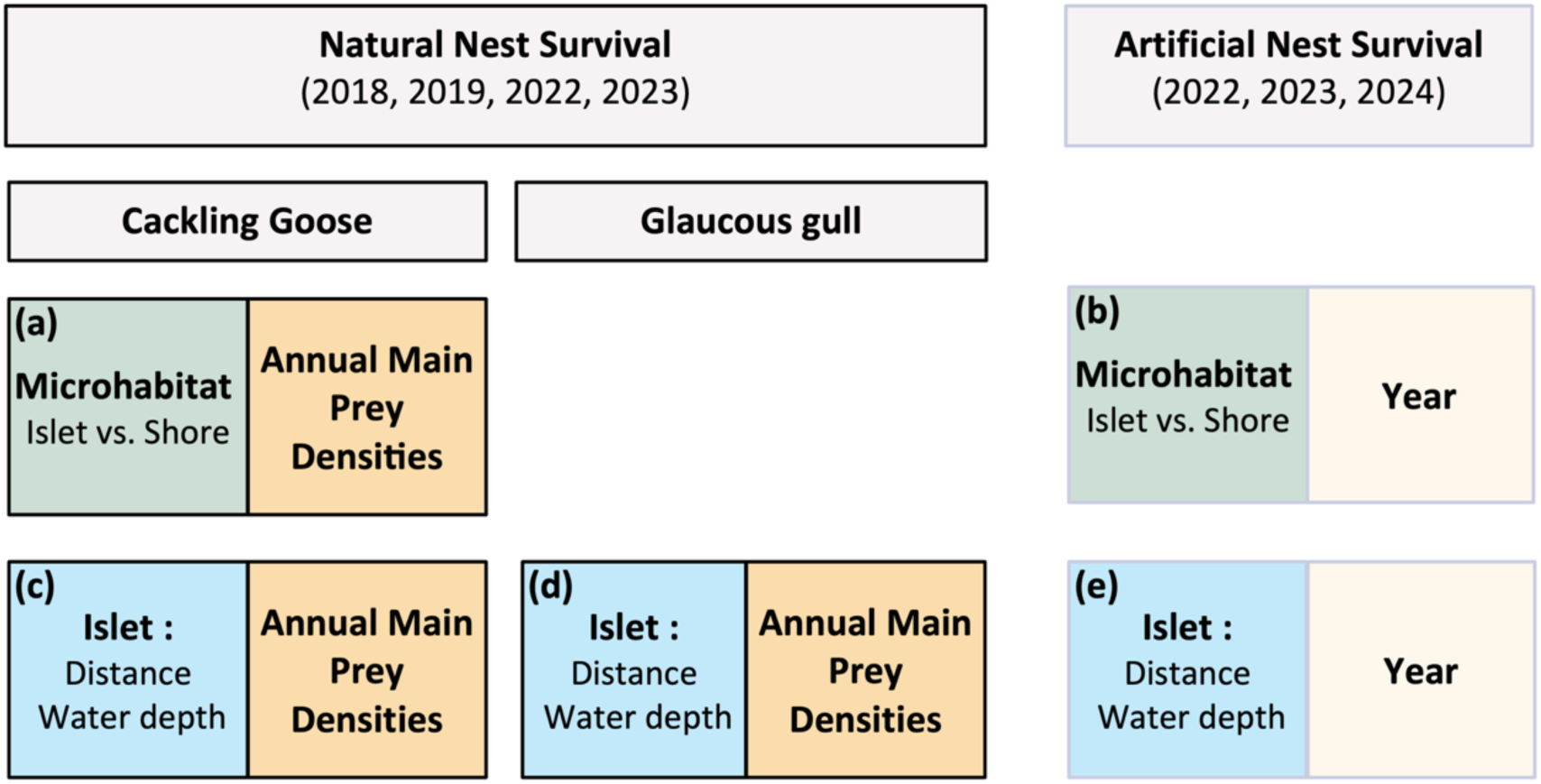
Design for assessing nest survival of natural and artificial nests in relation to nest microhabitat (a-b: islet vs. shore), or islet characteristics (c-e: distance to shore and water depth), and annual main prey densities (a,c,d: lemmings and snow goose nests) or interannual variation (e: year) in Bylot Island, Nunavut, Canada. Microhabitat effect could not be tested for glaucous gulls due to low sample size.

Moreover, we did not collect data in 2020 and 2021 due to the COVID-19 pandemic.

### Main prey availability

We estimated lemming density in 2018, 2019, 2022 and 2023 from live-trapping sessions conducted in the Qarliturvik Valley (73°08’N; 80°08W), 30 km north of the snow goose colony. Three sessions were conducted each summer (mid-June, mid-July and mid-August) in two 11-ha permanent grids, one in mesic tundra and one in wetland habitat. Each grid contained 144 Longworth traps, spaced 30 m apart. Sessions lasted 3 days, with traps checked every 12 hours. Captured lemmings were identified to species and marked with Passive Integrated Transponder tags before release. Using the mark-recapture method (see Fauteux, Gauthier, and Berteaux 2015), we estimated annual lemming density as the mean of June and July densities for each species and grid, a period which corresponds to the incubation period for both cackling goose and glaucous gull.

We estimated snow goose nest density in the study area in 2018, 2019, 2022, and 2023 following a multi-step process presented in Moisan et al. (2025). Within the goose colony (inside orange polygon Fig. 1c, d, e), whose outline was delimited annually using a GPS receiver aboard a helicopter, we assessed nest densities separately in wetlands and mesic habitats, as geese generally nest more densely in wetlands on Bylot Island (Lecomte, Gauthier, and Giroux 2008). First, wetlands were delineated through photointerpretation of high-resolution (∼30 cm) satellite images. Habitat not classified as freshwater or wetlands were considered mesic. Second, we calculated nest density in mesic habitat using systematic nest sampling and observations of breeding individuals, whereas wetlands densities relied solely on systematic nest sampling.

Third, we multiplied the annual wetland and mesic areas within the goose colony by their respective nest density to estimate the total annual nest count. Finally, we divided this count by the study area size (150 km^2^) to obtain the annual nest density for the entire study area.

In 2024, since the entire study site only became accessible in late July, we could not obtain these same estimates of lemming and snow goose nest densities. Lemming trapping was not conducted in June but in July and over smaller grids. For snow geese, a systematic nest search was carried out within a 0.2 km^2^ intensively studied core area at the center of the colony. This survey is conducted annually, typically during goose incubation, but in 2024 it was performed shortly after the goose hatching period as the site was inaccessible earlier in the season. For 2024, we inferred snow goose nest density from the remaining empty nest cups, providing a minimal estimate. This provided a basis for year-to-year comparison. We were thereby able to distinguish general differences in the availability of main prey between 2022, 2023 and 2024 (see Supporting information for annual main prey densities), during which we conducted artificial nest experiments.

### Cackling goose and glaucous gull nest monitoring

We conducted nest monitoring for glaucous gulls and cackling geese annually during the incubation period. We found 94% of nests monitored while systematically searching wetland patches in June and opportunistically discovered the remaining nests throughout the nesting period. We conducted nest searches by teams of 2-3 people equipped with binoculars, walking through known and potential nesting sites throughout the study area. Glaucous gull nests are conspicuous on elevated mounds, and individuals often begin mobbing from a considerable distance. Cackling goose nests can be harder to spot as females lay flat on their nests, making them more cryptic. However, they can display defensive behaviors and alarm calls when approached. Benefiting from the open landscape to directly spot nests and using parental behavior cues, we are confident that we found nearly all nests.

For each nest found, we recorded its GPS position, identified the species and noted the microhabitat type (islet or shore). To evaluate the influence of islet characteristics on nest survival, we measured, in the field, the shortest distance (±1m) from the islet to shore and the maximum depth (±5cm) along this distance. We floated eggs to estimate age and hatch date (Liebezeit et al. 2007) and checked nests opportunistically throughout the incubation to record signs of predation. We revisited nests on or shortly after their estimated hatch dates to determine their fate. We deemed a nest successful if at least one egg hatched (at least one membrane or gosling visible) or if the nest remained active (at least one warm egg) by the end of the monitoring period. We classified nests as unsuccessful if they were completely depredated (found empty or layered with fragments of eggshells with membrane still attached) during the monitoring period. We excluded a total of 39 out of 229 monitored nests from analyses because they were either: found depredated on the first visit (0.9%, n = 2), visited only once (6.1%, n = 14), abandoned by parents (0.9%, n = 2), had an uncertain fate (3.1%, n = 7), lacked a microhabitat identification (3.1%, n = 7), or had to be excluded for other reasons (e.g., predation by an avian predator due to the observer) (3.1%, n = 7).

### Artificial nest experiments

We conducted artificial nest experiments between June 28 and July 13, in 2022 and 2023, during the cackling goose and glaucous gull incubation period, and between July 21 and 31 2024, shortly after the bird hatching period. We deployed 71 experimental units in 2022, 105 in 2023 and 73 in 2024. Each experimental unit consisted of a triad of artificial nests placed in an equilateral triangle, 20 ± 3 meters apart: one artificial nest on an islet and two artificial nests on the nearest shore (see Supporting information for experimental design). The use of triads allowed us to test microhabitat effects while ensuring that each experimental unit was visited by a predator (see below). We placed artificial nests 20 meters apart, a distance at which arctic fox detection is generally high (Beardsell et al. 2021, Weiss-Blais 2025). To avoid defense of artificial nests by nearby nesting birds, we placed each nest >10m from red-throated loon, cackling goose, or snow goose nests (Robertson 1995; Bêty et al. 2001) and >50 m from glaucous gull nests (Burger and Beer 1975).

For each artificial nest, we formed a small depression in the ground, where we placed two standard chicken eggs on a bed of goose down, and inserted a colored nail in the down to help identify depredated nests. To limit detection by avian predators, we covered nests with lichen (genus *Bryoria* or *Gowardia),* completely concealing the eggs and down. We inserted a goose feather into the lichen cover to secure it over the nest. We handled all nest material using latex gloves to minimize human scent. We measured the distance between the shore and islet (±1m), as well as the greatest water depth associated with this distance (± 5cm). We revisited all nests in the triad at 24 ± 3-hour intervals and ended the experiment when at least one of the triad’s nests was depredated.

In summer 2022 and 2023, we conducted a parallel experiment using motion-triggered cameras to confirm whether arctic foxes were responsible for depredating covered artificial nests. We placed cameras ∼3 m from 40 single artificial nests deployed in wetland habitats in the study area. Arctic foxes depredated 33 nests and one raven (*Corvus corax*) depredated one, while six were not depredated, thus confirming that arctic foxes were the main predator of covered artificial nests in our study area.

To compare the survival of nests located in different microhabitats, knowing that an active predator, most likely a fox, came at ≤20m from each artificial nest, we excluded triads for which none of the two shore nests were depredated (6.4%, n = 16). This approach allowed us to reduce the likelihood that a nest remained non-depredated simply because the experimental site was not detected by a fox. We also excluded triads when the revisit interval was too long and predation occurred during that interval (1.6%, n = 4), or when the experiment was interrupted by the observer before any predation occurred (4.4%, n = 11). Although arctic foxes were the primary predators of artificial nests, avian predators could also prey upon them (see above). To test our hypothesis that nest survival would be higher for nests less accessible to foxes, we also excluded triads in which one or more artificial nests were depredated by avian predators, as indicated by the presence of eggshell remains near a depredated nest (Hall and Arnold 1962). While eggshell remains were found in only 6% of artificial nests, we excluded 15.7% of triads (n = 39) due to evidence of avian predation in at least one of the three nests.

### Statistical analyses

We tested whether nest survival of cackling goose nests i) differed between islet and shore nests, and ii) was affected by main prey densities (lemming and snow goose nest densities) (Fig. 3a). We used generalized linear mixed models with a binomial distribution (R package *lme4*, version 1.1.30 (Bates et al. 2015)) and the logistic-exposure method, which accounts for variations in nest monitoring length (i.e., exposure) and does not require assumptions about the timing of nest loss (Shaffer 2004). In this method, exposure is incorporated into the link function to estimate daily nest survival. We considered a nest to have survived (1) if at least one egg hatched or was still being incubated by a parent at the end of the monitoring; otherwise, it was deemed depredated (0). For successful nests, we defined exposure as the number of days from discovery to either the hatch date (observed or estimated) or to the last monitoring date if eggs were still incubating. For depredated nests, we calculated exposure as the number of days from discovery to the midpoint between the last active date and the date it was determined to be depredated, since the exact date of failure was not known. Our global model included nest microhabitat (islet or shore), lemming density and snow goose nest density as fixed effects, with the year as a random factor to control for other interannual variations. We tested the inclusion of nesting zones, identified via cluster analysis (see Duchesne et al. 2021), as random factor in the global model to account for potential spatial correlation. These zones had species-specific diameter and could be used annually for nesting. However, including this variable did not affect our results (not shown).

We tested whether survival of artificial nests i) differed between islet and shore nests, and ii) varied annually (Fig. 3b). Prior to analysis, we filtered the dataset by triad and removed one depredated shore nest per triad. When both shore nests were depredated, we removed one at random, and when only one was depredated, we removed that nest. This procedure retained a paired shore and islet nests, both of which could either have survived or been depredated. This design allowed us to directly compare nest survival across microhabitats, knowing that an active predator, most likely a fox, came at ≤20m from each artificial nest. We analyzed this dataset using logistic regression models (logit-link and binomial distribution). We classified an artificial nest as successful (1) when at least one egg remained in the nest, and as depredated (0) otherwise. Our global model included microhabitat (islet or shore), year and their interaction as fixed effects. Pair ID was included as a random effect to account for the paired experimental design.

We assessed whether nest survival of cackling goose and glaucous gull nests varied with islet distance to shore, water depth, and annual densities of main prey (lemmings and snow goose nests) (Fig. 3c, 3d). We used logistic-exposure models, aiming to include distance to shore, water depth, lemming density and snow goose nest density as fixed effects, with the year as a random factor in our global models. However, due to singular fit with a low number of groups in the random factor, we re-specified year as a fixed effect. Year was then highly correlated with prey densities, so it was removed. Multicollinearity between lemming and snow goose nest densities (VIF > 5) led us to create separate models for each variable. This multicollinearity was most likely due to the limited number of monitored years, as indicated by Pearson’s correlation coefficient: 0.80 with our four years of data compared to 0.18 when including 13 years (2010-2019, 2022 and 2023). Additionally, we removed the water depth variable for cackling geese due to convergence issues.

We assessed whether nest survival of artificial islet nests varied annually and with islet distance to shore and water depth (Fig. 3e). We analyzed these data using logistic regression models with logit-link, setting distance to shore, water depth, year and the interaction between distance and year (or water depth and year) as fixed effects in the global models.

When applicable, we tested the inclusion of islet ID as a random factor in the global models for all nest types, since a few islets were reused across the years. We found that adding this variable did not affect the overall results (not shown) and therefore did not include it in the global models. Given movement constraints on foxes (e.g., jump length and the need to swim in deep water), we suspected potential nonlinear effects of distance to shore and water depth (Corbeil-Robitaille et al. 2024). To investigate this, we applied distance-weighted transformations (*e^−α/Distance^* and *e^−α/Depth^*) to Euclidean distance and depth variables for all three nest types (Miguet, Fahrig, and Lavigne 2017). Following Carpenter et al. (2010), we tested various α values and used Akaike Information Criterion corrected for small sample size (AICc) to select the most parsimonious distance-weighted function for each variable, which were then used in the final model selection for each nest type (see Supporting information).

Across all analyses, we checked model assumptions, including linear relationship between logit(y) and each independent variable, multicollinearity, dispersion and checked for the presence of outliers (R packages *DHARMa*, version 0.4.6 (Hartig 2022); *car*, version 3.1.0 (Fox and Weisberg 2019) and blmeco, version 1.4 (Korner-Nievergelt et al. 2015)). From global models, we performed model selection by testing various combinations of variables, including the null model. We ranked models using R package *AICcmodavg*, version 2.3.1 (Mazerolle 2023). We considered models with ΔAICc ≤ 2 to be competitive (Anderson and Burnham 2002). We visualized results using coefficients from the best-supported model (i.e., the model with the highest weight) and obtained 95% confidence intervals using bootstrap. We performed all analyses in R version 4.2.2 (R Development Core Team 2022).

## Results

### Effects of refuge use on nest survival

We monitored 132 cackling goose nests in the study area across four breeding seasons (2018, 2019, 2022, and 2023), with 69 nests located on islets and 63 on shore (see yearly sample size in Supporting information). Nest survival was best explained by nest microhabitat and annual snow goose nest density. Specifically, we found moderate evidence of higher nest survival rates for nests on islets compared to those on shore, along with strong evidence of increased survival with higher annual snow goose nest density (Table 1, Fig. 4; see full model selection in Supporting information). At snow goose nest densities of 23 and 161 nests/km^2^ (corresponding, respectively, to the lowest and highest densities observed), nest survival rate was estimated to vary from 0.01 to 0.96 on shores and 0.27 to 0.99 on islets, based on an average exposure of 16 days. In contrast, we found little to no evidence of an effect of annual lemming density, with this variable even excluded from one of the competitive models (Table 1). When excluding data from 2022, the year with the lowest snow goose density, we still observed a moderate effect of nest microhabitat but found little to no evidence of an effect of annual snow goose nest density.

**Fig. 4.**
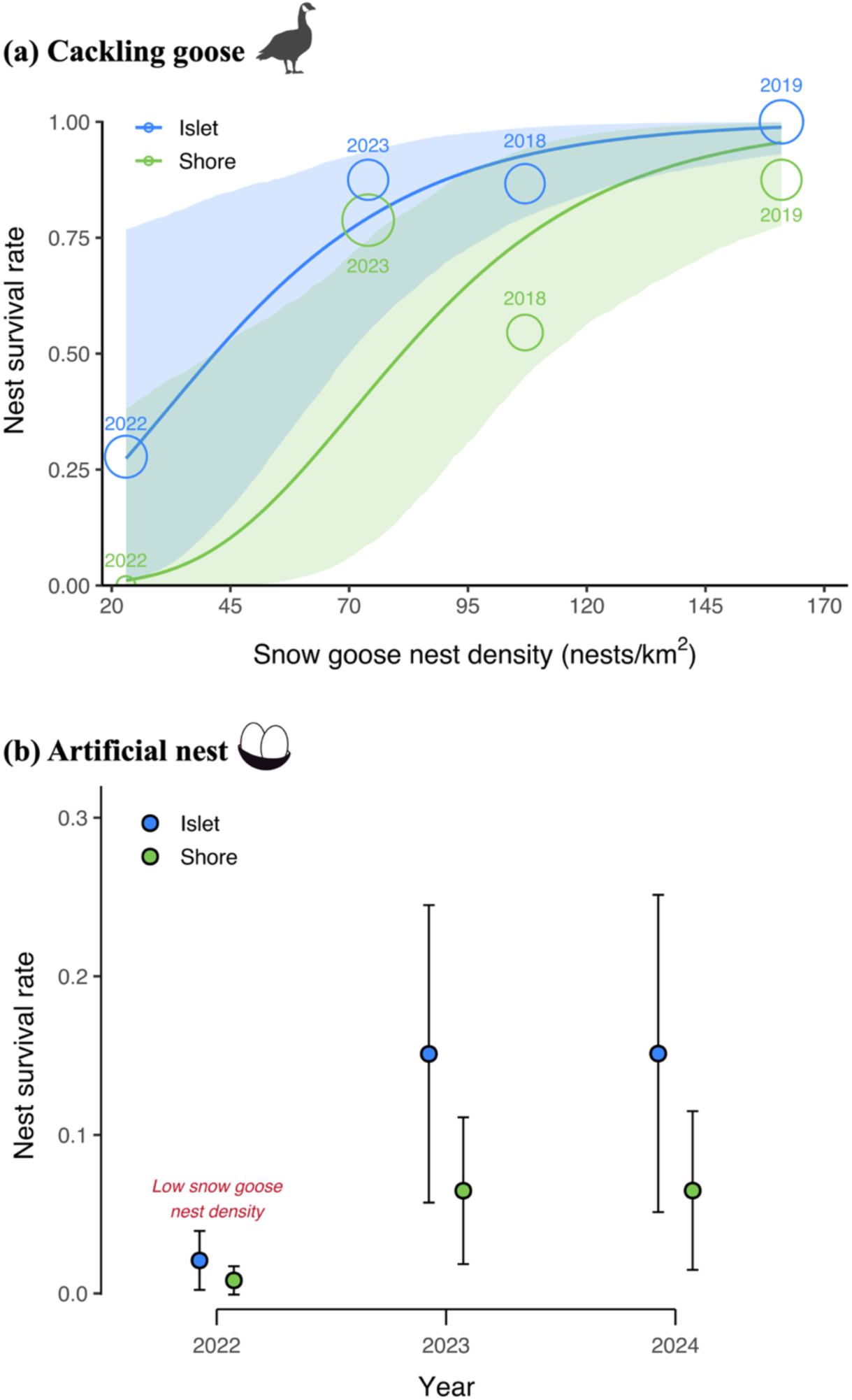
(a) Effect of nest microhabitat (islet vs. shore) and annual snow goose nest density on the survival rate of cackling goose nests (2018, 2019, 2022, 2023; N= 132). (b) Interannual variation in the survival rate of artificial nests across both microhabitats (2022-2024; N = 358). Year 2022 was characterized by exceptionally low snow goose nest density (see Methods). In both panels, solid lines (a) and dots (b) show mean model predictions over the average nest monitoring period (16 days in (a); 29 hours in (b)), with 95% confidence intervals (a) and standard errors (b). Color denotes nest microhabitat. Open circles sizes are proportional to the number of observed nests contributing to each survival estimate.

**Table 1.**
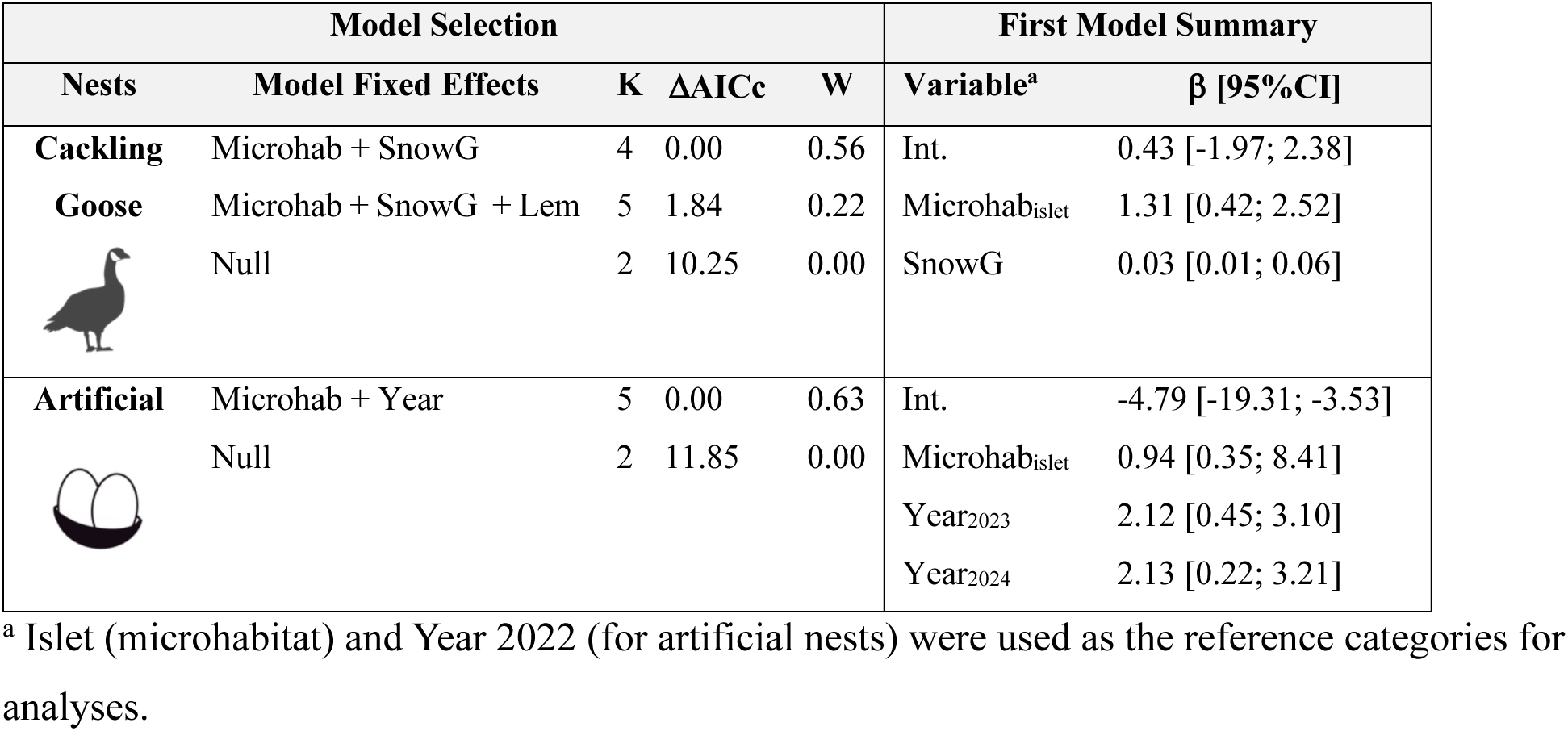
Generalized linear model selection of the effects of nest microhabitat (Microhab: islet vs. shore) and main prey densities (lemming (Lem) and snow goose nests (SnowG)) on probability of nest survival for cackling goose nests (2018, 2019, 2022, 2023) and artificial nests (2022-2024). Left panel reports null and competitive models (ΔAICc ≤ 2), with number of parameters (K), change in AICc from best-supported model (ΔAICc) and Akaike weights (W).The right panel reports estimated coefficients (β) with their 95% confidence interval (CI) of the model with the smallest AICc. Full model selection is presented in Supporting information.

| Model Selection |  |  |  |  | First Model Summary |  |
| --- | --- | --- | --- | --- | --- | --- |
| Nests | Model Fixed Effects | K | $\Delta AICc$ | W | Variable<sup>a</sup> | $\beta$ [95%CI] |
| **Cackling Goose** | Microhab + SnowG | 4 | 0.00 | 0.56 | Int. | 0.43 [-1.97; 2.38] |
|  | Microhab + SnowG + Lem | 5 | 1.84 | 0.22 | Microhab<sub>islet</sub> | 1.31 [0.42; 2.52] |
|  | Null | 2 | 10.25 | 0.00 | SnowG | 0.03 [0.01; 0.06] |
| **Artificial** | Microhab + Year | 5 | 0.00 | 0.63 | Int. | -4.79 [-19.31; -3.53] |
|  | Null | 2 | 11.85 | 0.00 | Microhab<sub>islet</sub> | 0.94 [0.35; 8.41] |
|  |  |  |  |  | Year<sub>2023</sub> | 2.12 [0.45; 3.10] |
|  |  |  |  |  | Year<sub>2024</sub> | 2.13 [0.22; 3.21] |

<sup>a</sup> Islet (microhabitat) and Year 2022 (for artificial nests) were used as the reference categories for analyses.

A total of 179 triads from the artificial nest experiments conducted in 2022, 2023, and 2024 were used for analysis. With a mean exposure time of 29 hours, the results of the artificial nest experiments thus approximate a daily nest survival rate. We found moderate evidence of a microhabitat effect on artificial nest survival, with survival rates on islets being 1-8% higher than on the shore, depending on the year. Additionally, we found strong evidence of a year effect, with nest survival rates of 0.11 in 2023 and 2024, compared to 0.01 in 2022, when snow goose nest density was exceptionally low (Table 1, Fig. 4; see full model selection in Supporting information).

### Effects of refuge quality on nest survival

We monitored natural nests located on islets with known physical characteristics (distance to shore and water depth) across four breeding seasons (2018, 2019, 2022, and 2023; see Supporting information for sample sizes). We monitored 67 cackling goose nests situated at a median distance of 9 m from shore (range: 1–45 m) and separated from the shore by a median maximum water depth of 30 cm (range: 3–40+ cm). We also monitored 55 glaucous gull nests located on islets at a median distance of 15 m from shore (range: 1–35 m) and separated from the shore by a median maximum water depth exceeding 40 cm (range: 7–40+ cm).

For natural nests, we found little to no evidence that distance to shore or water depth influenced nest survival in either species, with the later excluded from competitive models (Table 2, see selection of decay variables and full model selection in Supporting information). Still, the best-supported model suggested that nests located on islets further from shore presented higher nest survival rates, ranging from 0.75 to 1.00 for cackling goose (at 1 versus 45 meters from shore, based on average exposure of 16 days) and from 0.58 to 0.94 for glaucous gull (at 1 versus 35 meters from shore; based on average exposure of 15 days). However, as confidence intervals overlapped 0 for both species, these ranges should be interpreted with caution due to the limited evidence supporting an effect of distance. Note that water depth analysis was not feasible for cackling goose due to convergence issues, but an examination of raw data showed no strong survival pattern associated with either physical characteristic (Fig. 5). Additionally, survival of nests located on islets was strongly associated with annual snow goose nest density. As snow goose nest density increased from 23 to 161 nests/km^2^, nest survival rates rose from 0.62 to 1.00 for cackling geese and from 0.42 to 0.97 for glaucous gulls. In contrast, lemming density did not appear in any competitive models (Table 2, see full model selection in Supporting information).

**Fig. 5.**
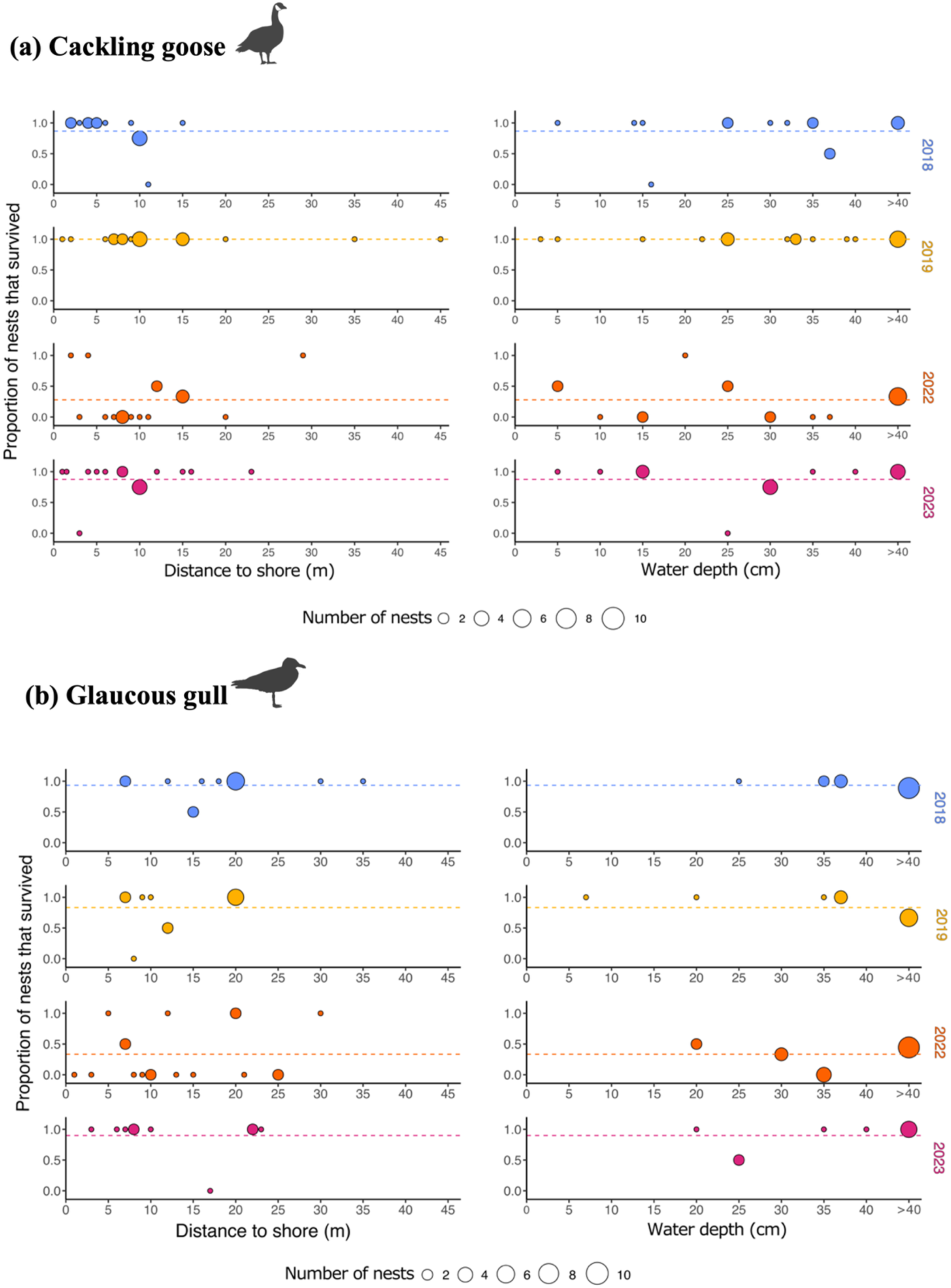
Proportion of nests that survived the monitoring interval, for (a) cackling geese and (b) glaucous gulls, on islets of varying distance from shore and water depth, presented by year (2018, 2019, 2022, and 2023). Points represent observed data, with size relative to the number of observations. Dashed lines indicate yearly observed mean nest survival.

**Table 2.**
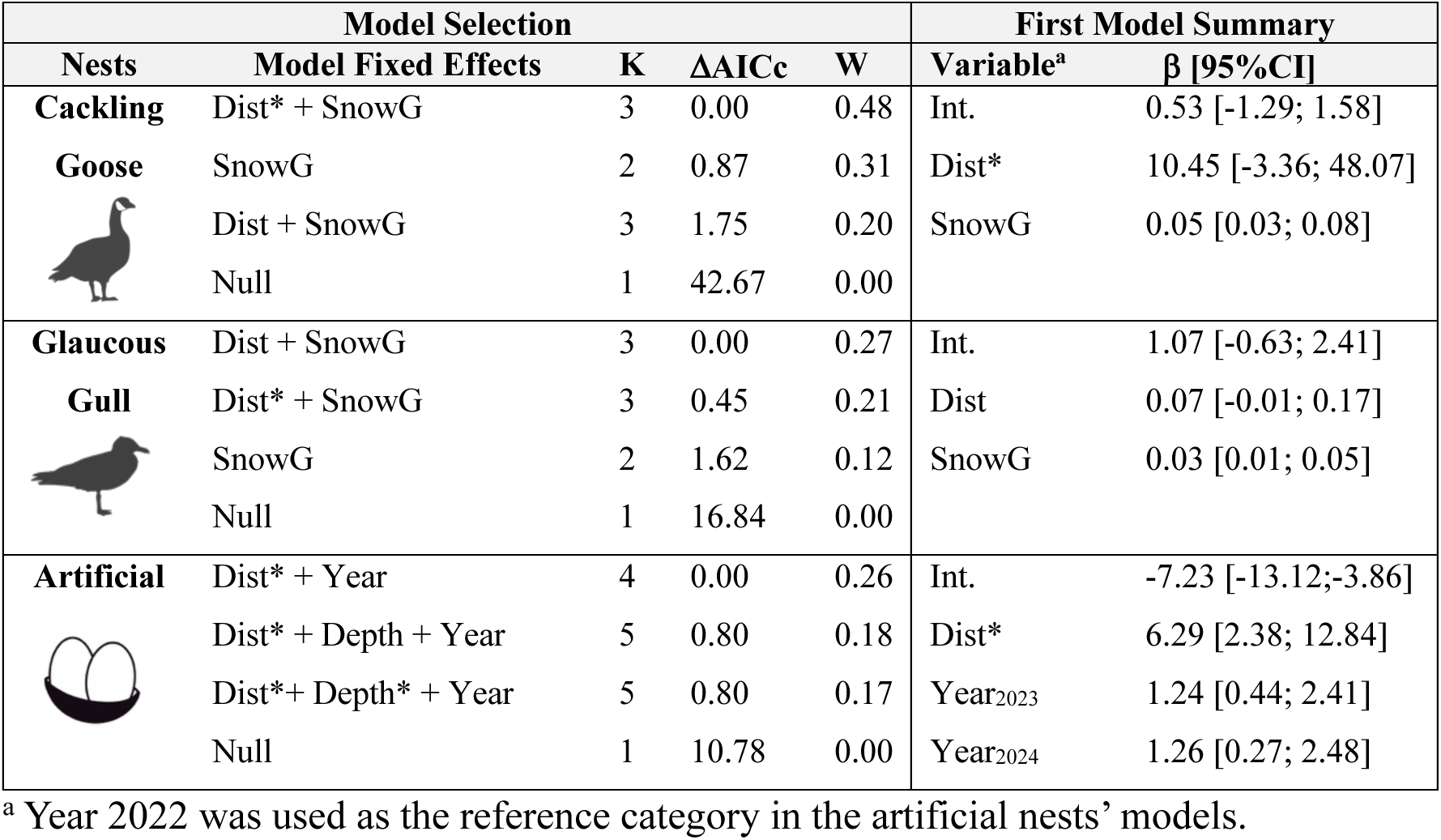
Generalized linear model selection of the effects of islet physical characteristics (distance to shore (Dist) and water depth (Depth)), and annual main prey densities (lemming and snow goose nests (SnowG)) or year, on the probability of nest survival for cackling goose nests (2018, 2019, 2022, 2023), glaucous gull nests (2018, 2019, 2022, 2023) and artificial nests (2022-2024). Star «*» indicates that a distance weighted function was used (see Methods). Left panel reports null and competitive models (ΔAICc ≤ 2), with number of parameters (K), change in AICc from best-supported model (ΔAICc), Akaike weights (W). The right panel reports estimated coefficients (β) with their 95% confidence interval (CI) of the model with the smallest AICc. Full model selection is presented in Supporting information.

| Model Selection |  |  |  |  | First Model Summary |  |
| --- | --- | --- | --- | --- | --- | --- |
| Nests | Model Fixed Effects | K | $\Delta AICc$ | W | Variable<sup>a</sup> | $\beta$ [95%CI] |
| **Cackling Goose** | Dist* + SnowG | 3 | 0.00 | 0.48 | Int. | 0.53 [-1.29; 1.58] |
|  | SnowG | 2 | 0.87 | 0.31 | Dist* | 10.45 [-3.36; 48.07] |
|  | Dist + SnowG | 3 | 1.75 | 0.20 | SnowG | 0.05 [0.03; 0.08] |
|  | Null | 1 | 42.67 | 0.00 |  |  |
| **Glaucous Gull** | Dist + SnowG | 3 | 0.00 | 0.27 | Int. | 1.07 [-0.63; 2.41] |
|  | Dist* + SnowG | 3 | 0.45 | 0.21 | Dist | 0.07 [-0.01; 0.17] |
|  | SnowG | 2 | 1.62 | 0.12 | SnowG | 0.03 [0.01; 0.05] |
|  | Null | 1 | 16.84 | 0.00 |  |  |
| **Artificial** | Dist* + Year | 4 | 0.00 | 0.26 | Int. | -7.23 [-13.12; -3.86] |
|  | Dist* + Depth + Year | 5 | 0.80 | 0.18 | Dist* | 6.29 [2.38; 12.84] |
|  | Dist* + Depth* + Year | 5 | 0.80 | 0.17 | Year<sub>2023</sub> | 1.24 [0.44; 2.41] |
|  | Null | 1 | 10.78 | 0.00 | Year<sub>2024</sub> | 1.26 [0.27; 2.48] |

<sup>a</sup> Year 2022 was used as the reference category in the artificial nests' models.

We monitored 179 artificial nests placed on islets over three breeding seasons on Bylot Island (2022, 2023, and 2024; see Supporting information for sample sizes). Nests were located on islets at a median distance of 7 m from shore (range: 1-20 m) and separated from the shore by a median maximum water depth of 25 cm (range: 5-40+ cm). Nest survival was best explained by distance to shore and year. Specifically, we found strong evidence for a nonlinear relationship between nest survival and distance to shore, as all competitive models included a distance-weighted function (Table 2). Nest survival on islets was lowest in 2022, characterized by exceptionally low snow goose nest density, and for nests located within a few meters from the shore. Nest survival increased sharply up to about 5 meters from the shore, then gradually stabilized (Fig. 6). For nests located 2 and 12 meters from shore (representing jumping distance and requiring entry into water, respectively), estimated nest survival rates, over a period of 29 hours, ranged from 0.03 to 0.19 in 2022 and 0.11 and 0.45 in 2023 and 2024. In contrast, there was little to no evidence of an effect of maximum water depth (whether on Euclidean or Decay scale) (see selection of decay variables and full model selection in Supporting information).

**Fig. 6.**
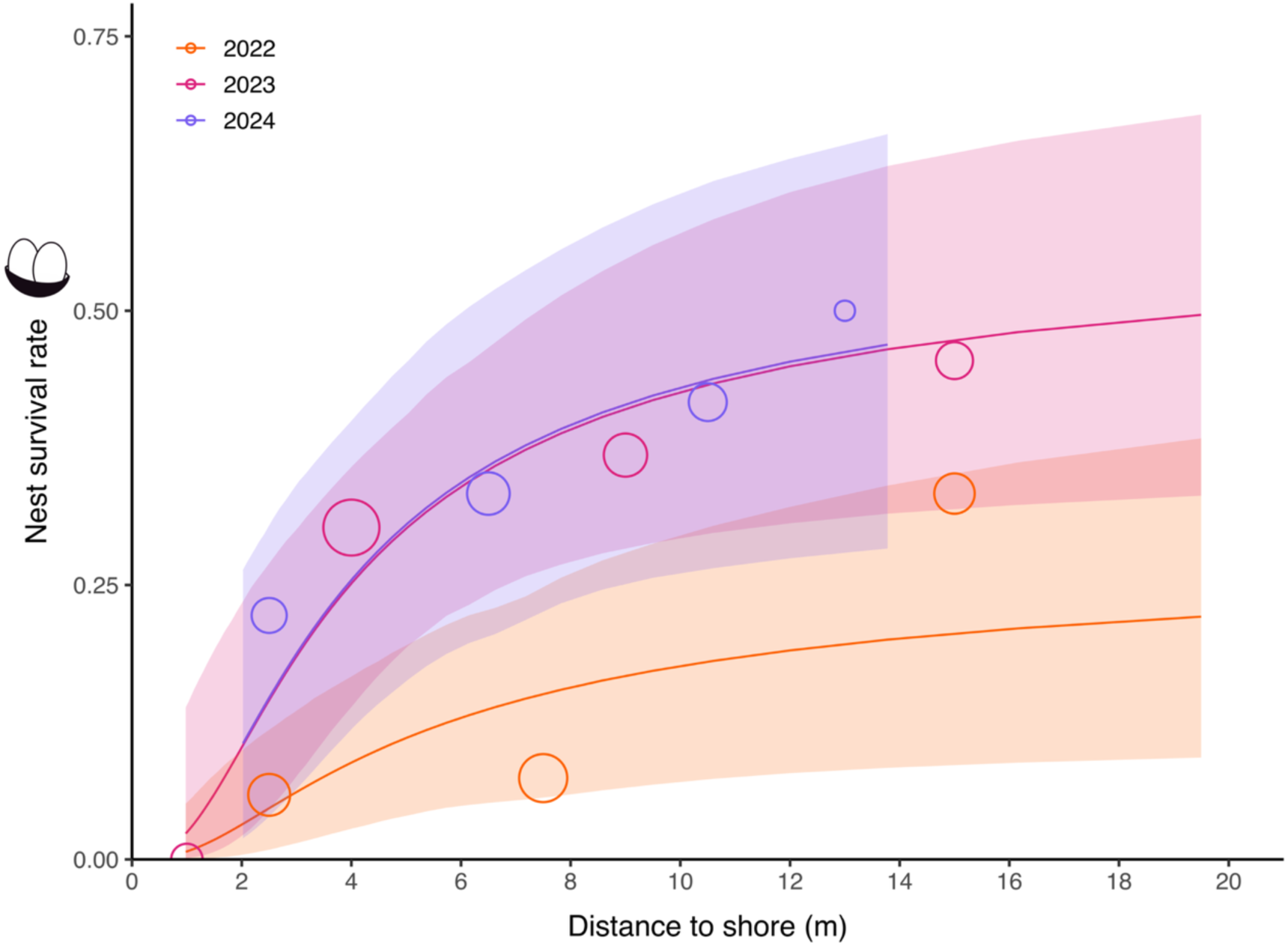
Interannual variation in the effect of distance to shore on nest survival rate of artificial nests located on islets (2022-2024). Circles represent observed data, with size proportional to the number of observations (N = 179). The full lines are the mean model prediction over the average nest monitoring period (29 hours) and are presented with their 95% confidence intervals.

## Discussion

Prey refuges can promote species occurrence and coexistence (Léandri-Breton and Bêty 2020; Duchesne et al. 2021). Using multi-year nest monitoring and field experiments, we showed that both natural and artificial nest survival was generally higher for nests located in microhabitats less accessible to the arctic fox, the main nest predator in the arctic tundra. As predicted (P1), survival was higher on islets than on shore. Among nests located on islets, survival was generally higher for those located at a greater distance from the shore. However, there was little evidence of a distance effect for cackling geese and glaucous gulls, both of which avoided nesting on islets near the shore (see below; Corbeil-Robitaille et al. 2024). No effect of water depth was detected for any nest types. Together, these results provide only partial support for P2. Though the use of islets reduced predation risk, our results also support the hypothesis that prey densities in the landscape can positively affect the survival of nests located on these partial refuges. This conclusion is mainly based on a single year characterized by exceptionally low density of one key prey species (snow goose nest density in 2022). The strong consistency in annual variation between natural and artificial nest survival patterns indicates that arctic foxes are more likely to target prey in less accessible microhabitats when their prey acquisition rate is reduced (see also Beardsell et al. 2025). As nest survival on islets within a snow goose colony appeared to be influenced primarily by snow goose nest density and less by lemming density, our results partially support P3. Overall, our study highlights the importance of considering both the physical landscape and prey densities to fully assess prey refuge quality and predator-prey interaction strengths (Fig. 7).

**Fig. 7.**
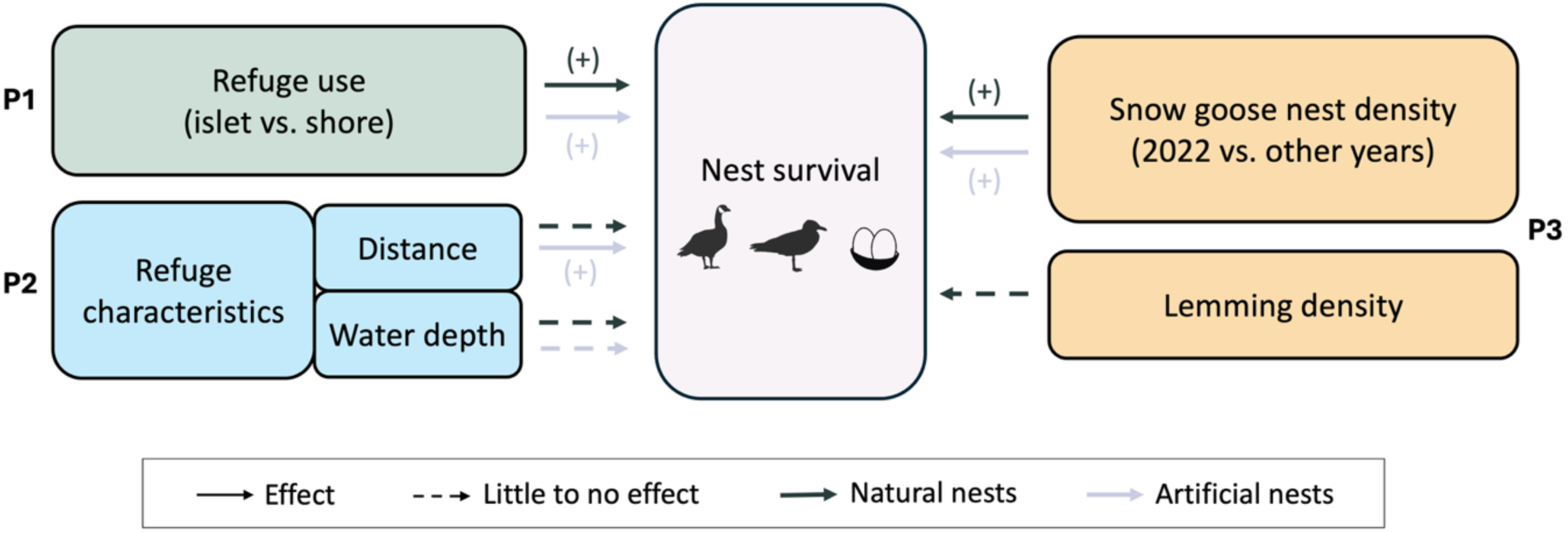
Schematic representation showing differences in the strength of evidence (solid vs. dashed lines) for biotic and abiotic drivers affecting natural (black) and artificial (grey) nest survival on pond and lake islets located within a snow goose colony on Bylot Island, Nunavut, Canada. Predictions are shown alongside the drivers (see Introduction).

### Pond and lake islets as partial refuges from arctic fox nest predation

As predicted, nest survival was higher on islets than on pond and lake shores for both natural and artificial nests. Since arctic foxes were the primary predators of artificial nests, our findings indicate that even small waterbodies (mean waterbody size of 0.008 km^2^; Corbeil-Robitaille et al. 2024) can hinder access to islets for these mammalian predators (see also Gauthier et al. 2015). The alignment between islet selection by birds (Eberl 1993; Bergman and Derksen 1977) and reduced nest predation risk supports an adaptative predator avoidance strategy, which could be particularly crucial to improve reproductive success in areas of high fox density (Clermont et al. 2021; Duchesne et al. 2021). Indeed, this strategy likely targets mammalian predators, as islets do not offer protection from avian predators.

These findings likely reflect physical and behavioral constraints on arctic foxes, which could reduce both the frequency and success of attacks on islets. Reaching islets may involve higher short-term costs, such as time involvement leading to missed foraging opportunities and increased energy expenditure from swimming (Alexander 2002), thermoregulation in cold water (Prestrud 1992), and cleaning or drying fur. Foxes may also avoid attacking nests on islets due to the higher risks of injury from harassing birds (e.g., to eye, skull and limbs), as limited movement in water may make it more difficult to anticipate and evade attacks (Mukherjee and Heithaus 2013; Shepard et al. 2013). Foxes, which typically rely on swift, agile charges to capture larger prey (Samelius and Alisauskas 2001), may find their mobility hindered in water.

This reduced speed can make it more challenging to evade bird parental defense and may also provide incubating birds with additional time to return to their nests if they were temporarily away, such as during an incubation recess. These factors likely contribute to fox hesitation to enter water, rendering islets into partial refuges for nesting birds.

Despite higher nest survival on islets, a significant number of cackling geese nested on pond and lake shores, even though most islets (>80%) remained unoccupied. One possible explanation is that late spring snow and ice cover reduced the availability of suitable nesting sites. During late spring break-up, some islets can be surrounded by an ice sheet at the beginning of the incubation period. However, cackling geese and glaucous gulls were rarely encountered nesting on islets surrounded by ice. This may result from risky conditions, as islets early in the season cannot be considered as refuges: a thick ice sheet may facilitate access for terrestrial predators, while later in the season, broken ice blocks could increase the risk of breeding failure due to shifting ice (Haynes et al. 2014). As cackling goose numbers have grown exponentially on Bylot Island since 1996 (Moisan et al. 2025), more individuals may be forced to nest outside islet refuges due to intra-specific competition and limited availability of suitable nesting sites.

### Subtle influence of islet physical characteristics on nest survival

The protective quality of islets against nest predation can vary with their physical characteristics. Nests on islets farther from shore generally exhibited higher survival rates, a pattern consistent with previous studies on island refuges (Strang 1976; Eberl 1993; Albrecht et al. 2006). In contrast, nests on islets close to shore faced greater predation risk from arctic foxes, especially those within a single jump reach, up to 4 meters (Bahr 1989). Beyond this distance, fox mobility is hindered by the presence of water, and the fitness costs of attacking a nest, particularly the risk of injury, may level off with increasing distance. While model selection identified distance to shore as a relevant predictor, its effect was little for natural nests but pronounced for artificial nests. This likely reflects the distribution of natural nests: over 70% of cackling goose nests and over 90% of glaucous gull nests were located more than 5 meters from shore. In contrast, artificial nest experiments indicate that the primary shift in predation risk occurs within the first 5 meters (Fig. 6). Consequently, because natural nests were mostly located beyond the 0-5 m zone where predation risk changes most sharply, our ability to detect this effect was likely reduced.

Alternatively, the limited effect of distance to shore observed for natural nests could result from the short islet-to-shore distances in the study area (maximum recorded distance: 54 meters). For example, Eberl (1993) found that red-throated loons which nest on islands >100 meters from shore were spared from fox predation, while those located <20 meters faced heavy losses. Similarly, Robertson (1995) reported that foxes avoided islands >50 meters from shore. The selection of islets farther from shore by cackling geese and glaucous gulls observed on Bylot Island (Corbeil-Robitaille et al. 2024) may still reflect an adaptative response to nest predation risk. However, under high predation pressure, this strategy may be less effective in the relatively small waterbodies of our study area compared to larger lake systems.

Apparent discrepancies between the results from natural and artificial nests regarding the effect of distance may also arise from a higher detection probability of natural nests by foxes. Indeed, foxes rely mainly on olfactory cues to detect covered artificial nests, while natural nests also provide visual cues (e.g., from parental activity; Martin, Scott, and Menge 2000).

Furthermore, odor-mediated detection varies with chemical composition (e.g., presence of down and incubating adult) and fluid dynamics (e.g., wind direction and strength) (Finelli et al. 2000). Odor cues from artificial nests on islets located farther from shore may dissipate more effectively, reducing detection and increasing nest survival. In contrast, natural nests may emit stronger cues, aiding detection by foxes even at greater distances. Strong distance effect in artificial nests could therefore arise from slightly lower detection and attack probabilities. However, the very limited number of natural nests found near the shore, and the resulting limited statistical power at short distances, most likely explains discrepancies in the effect of distance and suggests that birds avoid nesting on riskier islets located close to shore.

Water depth surrounding islets appeared less influential than distance to shore in determining islet protective quality, with little to no effect on natural and artificial nest survival. This contrasts with previous studies showing that greater water levels can reduce terrestrial predator access (Zoellick et al. 2004; Albrecht et al. 2006, Schmidt et al. 2023). This difference could be explained by limitations in the water depth variable used, which may not have accurately captured variation relevant to predator access. Other measurements, such as mean water depth along paths or intra-seasonal water depth variations, might better capture fox accessibility. However, the latter was irrelevant for artificial nests, as each experiment lasted a maximum of four days. Moreover, the composition of bottom sediments in small waterbodies could also influence predation risk. In shallow waterbodies presenting deep mud layers, foxes may face significant movement constraints, such as suction-induced drag (Liu, Huang, and Qian 2023), potentially exceeding the challenges of swimming. The joint effect of water and substrate barriers could therefore reduce fox efficacy and efficiency in depredating nests on islets, while increasing their vulnerability to bird counterattacks. Thus, water depth alone might be insufficient to predict fox accessibility.

This study focused on clutch predation risk but overlooked the potential threats to incubating adults. Unlike most avian predators, foxes pose a significant risk of injury or death to adults of both species. Cackling geese, for instance, exhibit short flushing distances (Kellett and Alisauskas 2011) and glaucous gulls leave one adult alone on the nest during alternating incubating and feeding shifts (Bustnes et al. 2001), making both species vulnerable to fox attacks. Islets farther from shore may enhance adult survival by reducing predator speed when approaching the nest and by providing earlier predator detection and more reaction time for incubating birds. Thus, selection for islet characteristics may not only be driven by short-term reproductive success but also by the need to support long-term adult survival, which may be even more critical. For example, pink-footed geese (*Anser brachyrhynchus*) in Svalbard show higher nesting success in slope habitats but still select cliff habitats, likely to enhance adult survival.

Fox-killed geese are often found in slope habitats but never in cliffs (Anderson et al. 2015). Similarly, in 2022, we observed a fox decapitate a female cackling goose incubating on a nest situated on a peninsula accessible to foxes, an observation that underscores the vulnerability of adults. Observational data on interactions between defensive nesting birds and offensive foxes would enhance our ability to model and quantify the strength of species interactions (Beardsell et al. 2025). Moreover, including incubating adult survival in future studies could reveal more complex drivers of nest site selection and population viability in natural communities (Rivers et al. 2025).

### Main prey availability as an indirect key factor in allospecific nest survival

Nest survival rates on islets and on pond and lake shores varied annually in both natural and artificial nests. While factors such as the physiological condition of incubating adults (Skinner et al. 1998) and their behavior (e.g., frequency and length of incubating recesses) may affect nest survival (Anderson et al. 2015), our experiments indicate that variation in predation pressure was a key driver of these interannual differences. Our conclusions are reinforced by the consistent patterns observed between natural and artificial nests, with survival of artificial nests being solely influenced by predation risk. Overall, our results strongly support the hypothesis that the protective quality of islets as prey refuges is modulated by predator foraging behavior, which appears closely tied to fluctuations in prey availability.

Predation pressure in prey partial refuges was closely linked to prey availability in our study area, with snow goose nest densities playing a dominant role. The summer of 2022 provided a rare opportunity of exceptionally low snow goose nest density within the colony (see Supporting information), coupled with low lemming density. This prey scarcity coincided with significantly lower nest survival rates on islets compared to other years. In periods of low prey abundance, foxes may experience a much lower energy acquisition rate, prompting increased foraging efforts (Beardsell et al. 2025). For instance, foxes are typically more active and travel longer daily distances when lemming densities are low (Beardsell et al. 2022). Such elevated activity could result in a greater time spent foraging in wetlands, increasing the likelihood of encountering and detecting nests on islets. Additionally, foxes may exhibit a greater willingness to take risks (Brown and Kotler 2004; Berger-Tal et al. 2009; Beardsell et al. 2025). Experiments with captive red foxes (*Vulpes vulpes*) have shown that hungry individuals spent more time foraging in riskier patches compared to when they were in better condition (Berger-Tal et al. 2009). Such behavioral changes, driven by energetic constraints, could explain an increased risk-taking behavior by foxes, even towards nests located in partial refuges such as islets.

We found strong evidence that snow goose nest densities in the landscape affected nest predation on islets, but little to no evidence that lemming densities did. The relatively low lemming densities throughout the study period, combined with the limited number of monitoring years and statistical constraints (i.e., multicollinearity inducing separate prey models; see Methods), may have reduced our ability to detect lemming effects. However, our findings align with prior observations of poor nest survival on islets during periods of prey scarcity. The presence of a large snow goose colony in our study area in most years likely buffered the impact of lemming fluctuations observed elsewhere (Iles et al. 2013; Flemming et al. 2019). Fox reproduction on Bylot Island is strongly tied to lemming availability (Giroux et al. 2012), yet when lemmings are scarce, only foxes with access to the goose colony reproduce, as it generally provides a reliable food source: eggs can be cached and consumed later, while goslings offer an immediate food supply for growing pups (Giroux et al. 2012). Remarkably, 2022 was the only year in the past two decades of monitoring with no fox reproduction observed across the entire ∼600 km^2^ study site of Bylot Island (Berteaux, unpublished data), emphasizing the critical role of goose colonies in mitigating food shortage when lemming densities are low. This buffering effect from the snow goose colony may explain the contrasting results previously reported on Bylot Island. Gauthier et al. (2015) and Beardsell et al. (2025), who investigated glaucous gull nest survival outside of the snow goose colony, showed that gull hatching success on both shore and islet nests was positively related to summer lemming density. Taken together, these observations highlight the importance of investigating how the energy intake rate influences fox behavior, rather than focusing solely on the influence of prey densities (Beardsell et al. 2025).

## Conclusion

Our study examined spatio-temporal variations in nest predation risk on pond and lake islets. We showed that islets serve as valuable partial refuges for some arctic-tundra nesting birds and that nest predation risk was primarily modulated by arctic fox foraging behavior in response to prey availability, rather than islet physical characteristics. We thereby emphasize the importance of considering both biotic and abiotic drivers of predation risk to untangle the effect of prey refuges on prey vulnerability and ability to persist in the landscape. In addition, enhancing our understanding of the fitness costs experienced by predators foraging in partial prey refuges could offer key insights into their behavioral flexibility and how it can shape community structure.

## Acknowledgements

This research was made possible thanks to the logistical support from the Bylot Island field station of the Center for Northern Studies located in Sirmilik National Park (Parks Canada) and Natural Resources Canada through the Polar Continental Shelf Program. We express our gratitude to the staff of Sirmilik National Park of Canada, the community of Mittimatalik and the Mittimatalik Hunters & Trappers Organization for their support of the Bylot Island long-term monitoring program. We also thank the many students and researchers who have contributed to data collection on Bylot Island over the years. We especially acknowledge Éléonore Douville, Sandrine Benoît, Matthieu Weiss-Blais, Eloi Gagnon, Thierry Grandmont and Louis Moisan, along with the fantastic goose and lemming teams for their fieldwork. We are sincerely grateful to Jeanne Clermont and Frédéric Dulude-de Broin for their insightful ideas and reflections, and to Marie-Christine Cadieux and Marie-Jeanne Rioux for their essential role in logistical support and data management. We also thank Alain Caron for his assistance with statistical analyses. We acknowledge the use of ChatGPT (Open AI) for assisting with code debugging and improvement, brainstorming for text organization, and enhancing readability and language. The authors take full responsibility for the final review and content of this publication. This study was funded by (alphabetical order): ArcticNet (a Network of Centres of Excellence Canada), the Canada Foundation for Innovation, the Canada Research Chairs Program, the Fonds de recherche du Québec−Nature et technologies (FRQNT), the Kenneth M. Molson Foundation, the Natural Sciences and Engineering Research Council of Canada (NSERC), the Northern Scientific Training Program (Polar Knowledge Canada), the Polar Continental Shelf Program (Natural Resources Canada) and the Weston Family Award for Northern Research through the Association of Canadian Universities for Northen Studies. M. Beaudoin received scholarships from NSERC and FRQNT.

## Ethics statement

Field techniques were approved by ethical committees of Université du Québec à Rimouski and Université Laval according to the Canadian Council on Animal Care (CCAC) guidelines. Field research was approved by the Joint Park Management Committee of Sirmilik National Park of Canada.

## Author contributions

M.B.: Conceptualization, Data curation, Formal analysis, Investigation, Visualization, Writing - original draft, Writing - review & editing;

A.B.: Formal analysis, Investigation, Supervision, Validation, Visualization, Writing - review & editing;

E.D.: Conceptualization, Formal analysis, Investigation, Methodology, Validation, Writing - review & editing;

M.-Z.C.-R.: Conceptualization, Formal analysis, Investigation, Methodology, Writing - review & editing;

P.L.: Funding acquisition, Investigation, Project administration, Resources, Validation, Writing - review & editing;

D.B.: Conceptualization, Funding acquisition, Methodology, Project administration, Supervision, Validation, Writing - review & editing;

J.B.: Conceptualization, Funding acquisition, Investigation, Methodology, Project administration, Resources, Supervision, Validation, Writing - review & editing.

## Data availability

Upon acceptance, data and code used for analyses will be made publicly available in an online repository.

