## Supplementary material for "Experimental and field evidence indicate that islet-nesting tundra birds experience reduced nest predation and benefit indirectly from high snow goose densities": Supplemental-Information.pdf

### Table of Contents:


A. Nests sample size and annual main prey densities ... 2

B. Snow goose nest densities from 2012-2023 ... 3

C. Design for covered artificial nest experiments ... 4

D. Distance-weighted functions ... 5

E. Full model selection for microhabitat and main prey densities, or year, effects on nest survival of cackling geese and artificial nests ... 7

F. Model selection for choice of decay physical characteristics variables for cackling goose, glaucous gull and artificial nest survival analysis ... 9

G. Full model selection for islet physical characteristics and main prey densities, or year, on nest survival of cackling goose, glaucous gull and artificial nest ... 13

References ... 19

### A. Nests sample size and annual main prey densities

Table A1. Number of natural and artificial nests monitored in different microhabitats (islet or shore), along annual summer lemming densities and snow goose nest densities in the study area, on Bylot Island, Canada (2018, 2019, 2022, 2024).

| Parameter | 2018 | | 2019 | | 2022 | | 2023 | | 2024 | |
|---|---|---|---|---|---|---|---|---|---|---|
| **Microhabitat** | Islet | Shore | Islet | Shore | Islet | Shore | Islet | Shore | Islet | Shore |
| **Number of nests** | | | | | | | | | | |
| Cackling goose | 15 | 11 | 20[a] | 16 | 18 | 3 | 16 | 33 | 0 | 0 |
| Glaucous gull | 15 | 1 | 12 | 0 | 18 | 0 | 10 | 2 | 0 | 0 |
| Artificial | 0 | 0 | 0 | 0 | 59 | 59 | 79 | 79 | 41 | 41 |
| **Summer lemming density** (ind/km$^2$) | 3.3 | | 233 | | 5.3 | | 4.7 | | ?[b] | |
| **Snow goose density** (nests/km$^2$) | 107 | | 161 | | 23 | | 74 | | ?[b] | |

[a] 18 out of 20 nests had known physical characteristics (distance and depth)

[b] See Table A2 for rough estimate

Table A2. Annual snow goose nest densities in the core of the study area, along July lemming densities in the study area, on Bylot Island, Canada (2022-2024).

| Main prey density | 2022 | 2023 | 2024 |
|---|---|---|---|
| **July lemming density** (ind/km$^2$) | 6.7 | 9.4 | 496.9 |
| **Snow goose nest density in the core of the colony** (nests/km$^2$) | 409 | 1056 | 814[a] |

[a] Inferred from a systemic nest survey conducted after the goose hatching period, which likely led to an underestimation of genuine nest density

Table A3. Sample size of artificial nests monitored on islets according to water depth and distance to shore in the study area, on Bylot Island, Canada (2022-2024)

| Distance / Depth | 1-4 m | | | 5-10 m | | | 11-15 m | | | 16-20 m | | |
|---|---|---|---|---|---|---|---|---|---|---|---|---|
| | 2022 | 2023 | 2024 | 2022 | 2023 | 2024 | 2022 | 2023 | 2024 | 2022 | 2023 | 2024 |
| **5-15 cm** | 6 | 11 | 1 | 4 | 7 | 2 | - | 1 | - | - | - | - |
| **20-30 cm** | 4 | 11 | 3 | 19 | 25 | 14 | 3 | 7 | 3 | - | - | - |
| **>30 cm** | 2 | 5 | 5 | 14 | 9 | 11 | 6 | 1 | 2 | 1 | 2 | - |

**B. Snow goose nest densities from 2012-2023**

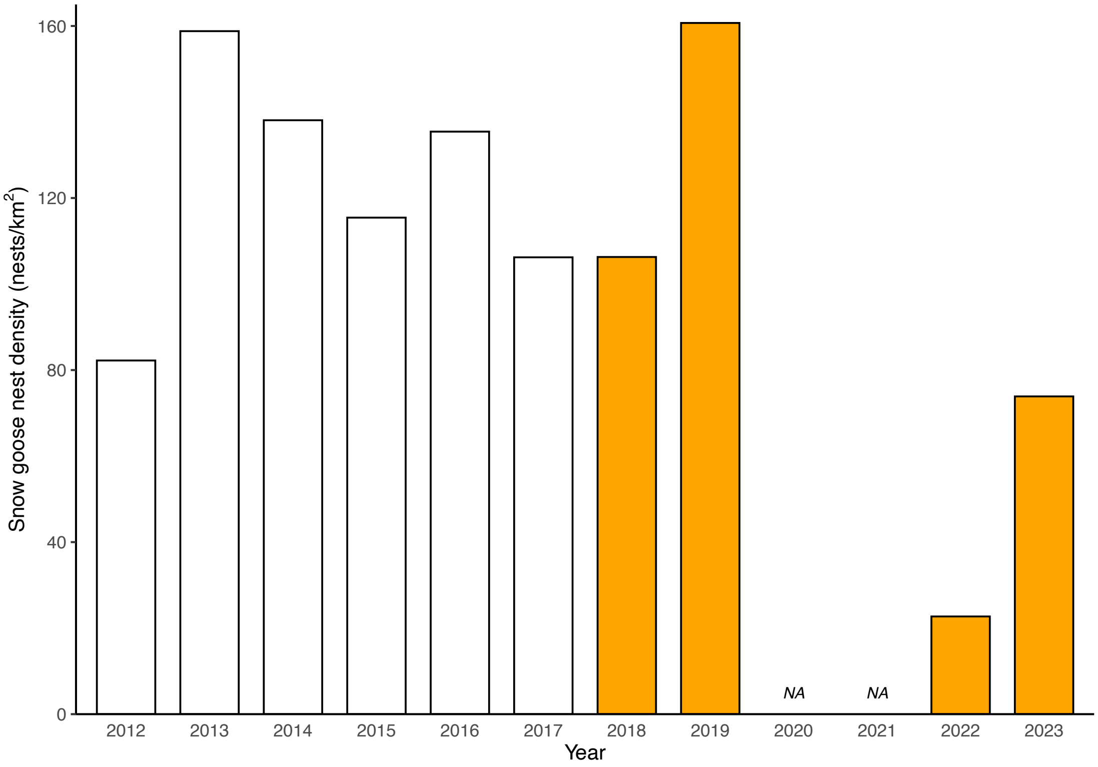


Figure B1. Snow goose nest densities (nests/$km^2$) in the 150 $km^2$ study area in Bylot Island over the past decade. Orange bars indicate data used in this study. No data are available for 2020-2021 due to COVID-19 pandemic.

**C. Design for covered artificial nest experiments**

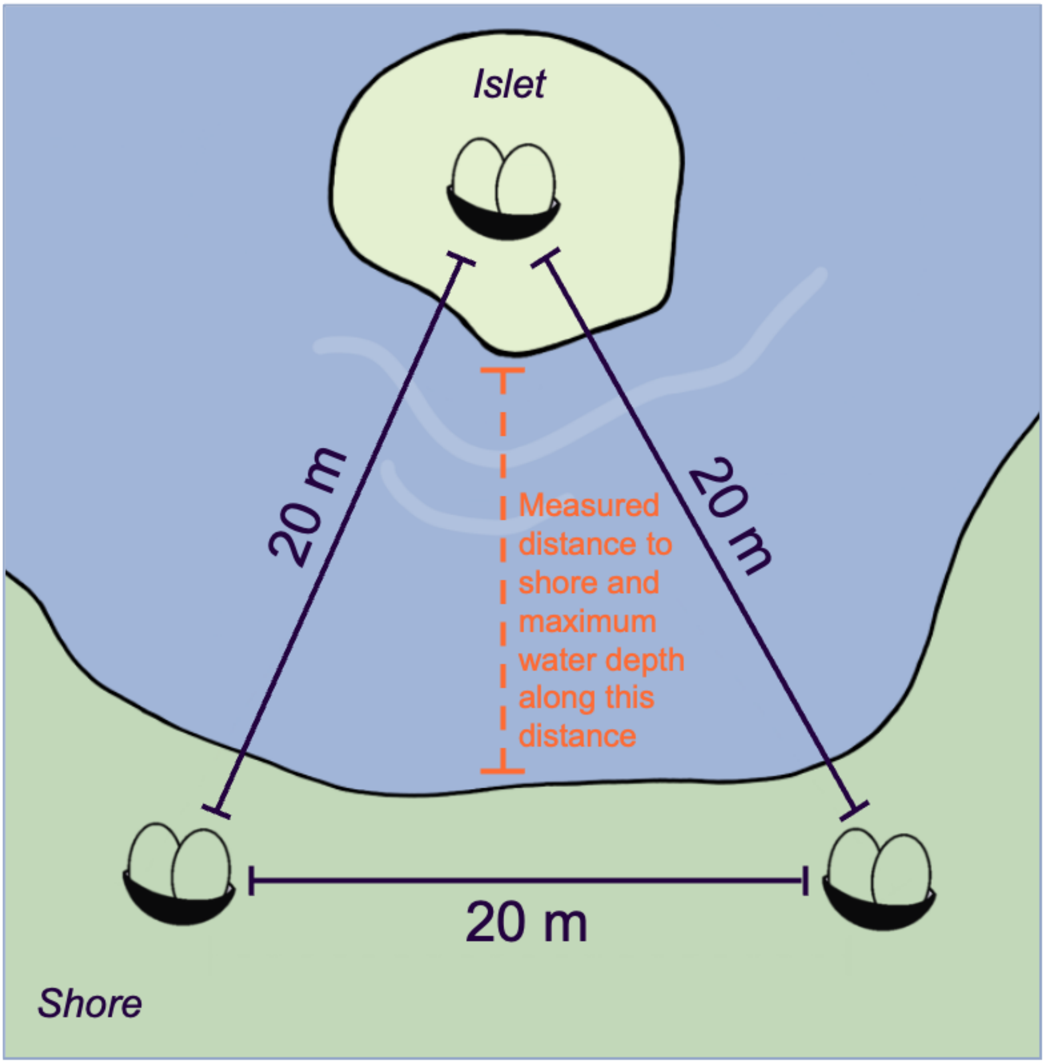


Figure C1. Layout of the experimental design: three paired covered artificial nests were deployed in triads in wetland habitats within the study area. The artificial nests were spaced 20 meters apart, with one nest located on an islet and the other two positioned on the nearest shore. Nests were covered with lichen to conceal the eggs and down. We measured the shortest distance from the islet to the shore, as well as the maximum water depth along this distance. This design allowed us to test microhabitat effects knowing that a fox came at ≤20m from each artificial nest.

**D. Distance-weighted functions**

The arctic fox can jump distances of up to 4 meters (Bahr 1989) and must swim in water deeper than 30 cm (based on measurements from 34 fox carcasses, M. Beaudoin, unpublished). Given these physical limitations, we applied distance-weighted functions to account for a potential decrease in the influence of distance to shore and water depth as the distance from the biological response point increases (Miguet, Fahrig, and Lavigne 2017).

For each Euclidean variable, we created decay variables using the negative exponential function $e^{-\alpha/Distance}$ or $e^{-\alpha/Depth}$, where α ranged between the minimum and maximum Euclidean values of distance or depth for each dataset. For cackling goose nests, we tested α values of 1, 5, 10, 20 and 45 for distance, but none for depth, as models failed to converge with depth variable. For glaucous gull nests, we tested α values of 1, 5, 10, 20, 35 for distance, and 7, 15, 30, and 41 for depth. For artificial nests, we tested α values of 1, 5, 10 and 20 for distance and 5, 15, 30, 60, and 75 for depth. These decay variables were scaled between 0 and 1, with higher values corresponding to the influence at great distance or depth.

For each nest dataset, we performed a separate model selection for each physical characteristic to identify which distance and depth decay variables best fit the data. For distance, we created a set of global models, each including the variables of interest (all variables from global model except distance) alongside one of the distance decay variables. For depth, we created similar models, each including the variables of interest (all variables from global model except depth) alongside one of the depth decay variables. Models were ranked using Akaike Information Criterion corrected for small sample size (AICc), ΔAICc and AICc-weights, using the R package *AICcmodavg*, version 2.3.1 (Mazerolle 2023). Models with ΔAICc ≤ 2 were considered competitive (Anderson and Burnham 2002). When more than one model was competitive, we chose the model with the best goodness of fit, indicated by the highest weight.

As we developed two global models for cackling geese and glaucous gulls (one per main prey), we conducted a sensitivity analysis comparing model selection outcomes for distance and depth decay variables across the global models. For glaucous gulls, we obtained the same selected distance and depth decay variables. However, the best-supported model differed for cackling geese, leading us to retain two distance decay variables for this specie.

For cackling goose nests, we retained *Distance decay* = $e^{-45/Distance}$ (from snow goose global model) and $e^{-1/Distance}$ (from lemming global model); for glaucous gull nests, *Distance decay* = $e^{-1/Distance}$ and *Depth decay* = $e^{-7/Depth}$; and for artificial nests, *Distance decay* = $e^{-1/Distance}$ and *Depth decay* = $e^{-75/Depth}$. These selected decay variables were then used in the final model selection, performed using different combinations of explanatory variables for each dataset.

**E. Full model selection for microhabitat and main prey densities, or year, effects on nest survival of cackling geese and artificial nests**

**Cackling goose nests**

Table E1. Generalized linear model selection of the effects of nest microhabitat (Microhab: islet vs. shore) and main prey densities (lemmings (Lem) and snow goose nests (SnowG)) on the probability of nest survival for cackling goose (2018, 2019, 2022 and 2023) on Bylot Island (N = 132). All candidate models are presented with their number of parameters (K), the change in AICc from the best-supported model (∆AICc), Akaike weight (W), and Log-Likelihood (LogLik). Competitive models (∆AICc < 2) are illustrated in bold.

| Models | K | AICc | ∆AICc | W | LogLik |
|---|---|---|---|---|---|
| **Microhab + SnowG** | **4** | **184.16** | **0.00** | **0.56** | **-87.92** |
| **Microhab + SnowG + Lem** | **5** | **186.00** | **1.84** | **0.22** | **-87.76** |
| Microhab | 3 | 188.17 | 4.01 | 0.08 | -90.99 |
| Microhab + Lem | 4 | 188.36 | 4.20 | 0.07 | -90.02 |
| SnowG | 3 | 189.03 | 4.87 | 0.05 | -91.42 |
| SnowG + Lem | 4 | 191.06 | 6.90 | 0.02 | -91.37 |
| Lem | 3 | 193.87 | 9.71 | 0.00 | -93.84 |
| Null | 2 | 194.41 | 10.25 | 0.00 | -95.16 |

Table E2. Coefficient estimates of competitive models from the generalized linear model selection of the effects of nest microhabitat (Microhab: islet vs. shore) and main prey densities (lemmings (Lem) and snow goose nests (SnowG)) on the probability of nest survival for cackling goose (2018, 2019, 2022 and 2023) on Bylot Island (N = 132). Coefficients' 95% confidence intervals are shown between square brackets.

| Models | Int. | $\text{Microhab}_{\text{islet}}$ | SnowG | Lem |
|---|---|---|---|---|
| Microhab + SnowG | 0.43 [-1.97;2.38] | 1.31 [0.42;2.52] | 0.03 [0.01;0.06] | |
| Microhab + SnowG + Lem | 0.10 [-2.49;2.23] | 1.34 [0.47;2.53] | 0.04 [0.01;0.08] | -0.004 [-0.02;0.02] |

### Artificial nests

Table E3. Generalized linear model selection of the effects of nest microhabitat (Microhab: islet vs. shore) and year on nest survival for artificial nests (2022-2024) on Bylot Island (N = 358). All candidate models are presented with their number of parameters (K), the change in AICc from the best-supported model (ΔAICc), Akaike weight (W), and Log-Likelihood (LogLik). Competitive models (ΔAICc < 2) are illustrated in bold.

| Models | K | AICc | ΔAICc | W | LogLik |
|---|---|---|---|---|---|
| **Microhab + Year** | **5** | **346.28** | **0.00** | **0.63** | **-168.05** |
| Microhab | 3 | 348.34 | 2.07 | 0.22 | -171.14 |
| Microhab + Year + Microhab:Year | 7 | 350.24 | 3.97 | 0.09 | -167.96 |
| Year | 4 | 350.93 | 4.66 | 0.06 | -171.41 |
| Null | 2 | 358.12 | 11.85 | 0.00 | -177.04 |

Table E4. Coefficient estimates of the unique competitive model from the generalized linear model selection of the effects of nest microhabitat (Microhab: islet vs. shore) and year on nest survival for artificial nests (2022-2024) on Bylot Island (N = 358). Coefficients' 95% confidence intervals are shown between square brackets.

| Models | Int. | $\text{Microhab}_{\text{islet}}$ | $\text{Year}_{2023}$ | $\text{Year}_{2024}$ |
|---|---|---|---|---|
| Microhab + Year | -4.79 [-19.31; -3.53] | 0.94 [0.35;8.41] | 2.12 [0.45;3.10] | 2.13 [0.22;3.21] |

### F. Model selection for choice of decay physical characteristics variables for cackling goose, glaucous gull and artificial nest survival analysis

#### Cackling goose nests

Table F1. Generalized linear model selection for the choice of the distance decay variable for analyzing the effects of distance to shore (Dist) and snow goose nest densities (SnowG) on the probability of nest survival for cackling geese nesting on islets (2018, 2019, 2022 and 2023) on Bylot Island (N = 67). The exponents in the Dist variables indicate the transformations tested. All candidate models are presented with their number of parameters (K), the change in AICc from the best-supported model (ΔAICc), Akaike weight (W), and Log-Likelihood (LogLik). Competitive models (ΔAICc < 2) are illustrated in bold.

| Models | K | AICc | ΔAICc | W | LogLik |
|---|---|---|---|---|---|
| **$Dist^{45}$ + SnowG** | **3** | **80.55** | **0.00** | **0.47** | **-37.09** |
| **$Dist^{20}$ + SnowG** | **3** | **82.22** | **1.67** | **0.20** | **-37.92** |
| $Dist^{10}$ + SnowG | 3 | 83.28 | 2.73 | 0.12 | -38.45 |
| $Dist^{1}$ + SnowG | 3 | 83.45 | 2.90 | 0.11 | -38.54 |
| $Dist^{5}$ + SnowG | 3 | 83.60 | 3.04 | 0.10 | -38.61 |

Table F2. Generalized linear model selection for the choice of the distance decay variable for analyzing the effects of distance to shore (Dist) and lemming densities (Lem) on the probability of nest survival for cackling geese nesting on islets (2018, 2019, 2022 and 2023) on Bylot Island (N = 67). The exponents in the Dist variables indicate the transformations tested. All candidate models are presented with their number of parameters (K), the change in AICc from the best-supported model (ΔAICc), Akaike weight (W), and Log-Likelihood (LogLik). Competitive models (ΔAICc < 2) are illustrated in bold.

| Models | K | AICc | ΔAICc | W | LogLik |
|---|---|---|---|---|---|
| **$Dist^{1}$ + Lem** | **3** | **109.00** | **0.00** | **0.41** | **-51.31** |
| **$Dist^{5}$ + Lem** | **3** | **109.98** | **0.98** | **0.25** | **-51.80** |
| **$Dist^{10}$ + Lem** | 3 | **110.91** | **1.92** | **0.16** | **-52.27** |
| $Dist^{20}$ + Lem | 3 | 111.95 | 2.96 | 0.09 | -52.79 |
| $Dist^{45}$ + Lem | 3 | 112.34 | 3.35 | 0.08 | -52.98 |

### Glaucous gull nests

Table F3. Generalized linear model selection for the choice of the distance decay variable for analyzing the effects of distance to shore (Dist), water depth (Depth) and snow goose nest densities (SnowG) on the probability of nest survival for glaucous gulls nesting on islets (2018, 2019, 2022 and 2023) on Bylot Island (N = 55). The exponents in the Dist variables indicate the transformations tested. All candidate models are presented with their number of parameters (K), the change in AICc from the best-supported model (ΔAICc), Akaike weight (W), and Log-Likelihood (LogLik). Competitive models (ΔAICc < 2) are illustrated in bold.

| Models | K | AICc | ΔAICc | W | LogLik |
|---|---|---|---|---|---|
| **$Dist^{35}$ + Depth + SnowG** | **4** | **79.15** | **0.00** | **0.24** | **-35.18** |
| **$Dist^{1}$ + Depth + SnowG** | **4** | **79.39** | **0.24** | **0.21** | **-35.30** |
| **$Dist^{20}$ + Depth + SnowG** | **4** | **79.63** | **0.48** | **0.19** | **-35.41** |
| **$Dist^{5}$ + Depth + SnowG** | **4** | **79.67** | **0.52** | **0.19** | **-35.44** |
| **$Dist^{10}$ + Depth + SnowG** | **4** | **79.84** | **0.69** | **0.17** | **-35.52** |

Table F4. Generalized linear model selection for the choice of the distance decay variable for analyzing the effects of distance to shore (Dist), water depth (Depth) and lemming densities (Lem) on the probability of nest survival for glaucous gulls nesting on islets (2018, 2019, 2022 and 2023) on Bylot Island (N = 55). The exponents in the Dist variables indicate the transformations tested. All candidate models are presented with their number of parameters (K), the change in AICc from the best-supported model (ΔAICc), Akaike weight (W), and Log-Likelihood (LogLik). Competitive models (ΔAICc < 2) are illustrated in bold.

| Models | K | AICc | ΔAICc | W | LogLik |
|---|---|---|---|---|---|
| **$Dist^{1}$ + Depth + Lem** | **4** | **93.38** | **0.00** | **0.32** | **-42.29** |
| **$Dist^{5}$ + Depth + Lem** | **4** | **94.31** | **0.93** | **0.20** | **-42.76** |
| **$Dist^{35}$ + Depth + Lem** | **4** | **94.57** | **1.19** | **0.18** | **-42.89** |
| **$Dist^{10}$ + Depth + Lem** | **4** | **94.85** | **1.47** | **0.15** | **-43.03** |
| **$Dist^{20}$ + Depth + Lem** | **4** | **94.88** | **1.50** | **0.15** | **-43.04** |

Table F5. Generalized linear model selection for the choice of the depth decay variable for analyzing the effects of distance to shore (Dist), water depth (Depth) and snow goose nest densities (SnowG) on the probability of nest survival for glaucous gulls nesting on islets (2018, 2019, 2022 and 2023) on Bylot Island (N = 55). The exponents in the Depth variables indicate the transformations tested. All candidate models are presented with their number of parameters (K), the change in AICc from the best-supported model (ΔAICc), Akaike weight (W), and Log-Likelihood (LogLik). Competitive models (ΔAICc < 2) are illustrated in bold.

| Models | K | AICc | ΔAICc | W | LogLik |
|---|---|---|---|---|---|
| **Dist + $Depth^{7}$ + SnowG** | **4** | **78.89** | **0.00** | **0.26** | **-35.05** |
| **Dist + $Depth^{15}$ + SnowG** | **4** | **78.92** | **0.03** | **0.25** | **-35.06** |
| **Dist + $Depth^{30}$ + SnowG** | **4** | **78.97** | **0.08** | **0.25** | **-35.09** |
| **Dist + $Depth^{41}$ + SnowG** | **4** | **79.00** | **0.10** | **0.24** | **-35.10** |

Table F6. Generalized linear model selection for the choice of the depth decay variable for analyzing the effects of distance to shore (Dist), water depth (Depth) and lemming densities (Lem) on the probability of nest survival for glaucous gulls nesting on islets (2018, 2019, 2022 and 2023) on Bylot Island (N = 55). The exponents in the Depth variables indicate the transformations tested. All candidate models are presented with their number of parameters (K), the change in AICc from the best-supported model (ΔAICc), Akaike weight (W), and Log-Likelihood (LogLik). Competitive models (ΔAICc < 2) are illustrated in bold.

| Models | K | AICc | ΔAICc | W | LogLik |
|---|---|---|---|---|---|
| **Dist + $Depth^{7}$ + Lem** | **4** | **94.21** | **0.00** | **0.25** | **-42.71** |
| **Dist + $Depth^{15}$ + Lem** | **4** | **94.25** | **0.03** | **0.25** | **-42.72** |
| **Dist + $Depth^{41}$ + Lem** | **4** | **94.25** | **0.04** | **0.25** | **-42.73** |
| **Dist + $Depth^{30}$ + Lem** | **4** | **94.26** | **0.05** | **0.25** | **-42.73** |

### Artificial nests

Table F7. Generalized linear model selection for the choice of the distance decay variable for analyzing the effects of distance to shore (Dist), water depth (Depth) and year on the probability of nest survival for artificial nests located on islets (2022-2024) on Bylot Island (N = 179). The exponents in the Dist variables indicate the transformations tested. All candidate models are presented with their number of parameters (K), the change in AICc from the best-supported model (ΔAICc), Akaike weight (W), and Log-Likelihood (LogLik). Competitive models (ΔAICc < 2) are illustrated in bold.

| **Models** | **K** | **AICc** | **ΔAICc** | **W** | **LogLik** |
|---|---|---|---|---|---|
| **$Dist^{1}$+ Depth + Year** | **5** | **198.15** | **0.00** | **0.43** | **-93.90** |
| **$Dist^{5}$+ Depth + Year** | **5** | **198.89** | **0.74** | **0.29** | **-94.27** |
| **$Dist^{10}$+ Depth + Year** | **5** | **199.66** | **1.51** | **0.20** | **-94.66** |
| $Dist^{20}$+ Depth + Year | 5 | 201.57 | 3.42 | 0.08 | -95.61 |

Table F8. Generalized linear model selection for the choice of the depth decay variable for analyzing the effects of distance to shore (Dist), water depth (Depth) and year on the probability of nest survival for artificial nests located on islets (2022-2024) on Bylot Island (N = 179). The exponents in the Depth variables indicate the transformations tested. All candidate models are presented with their number of parameters (K), the change in AICc from the best-supported model (ΔAICc), Akaike weight (W), and Log-Likelihood (LogLik). Competitive models (ΔAICc < 2) are illustrated in bold.

| **Models** | **K** | **AICc** | **ΔAICc** | **W** | **LogLik** |
|---|---|---|---|---|---|
| **Dist + $Depth^{75}$ + Year** | **5** | **200.58** | **0.00** | **0.26** | **-95.12** |
| **Dist + $Depth^{60}$ + Year** | **5** | **200.76** | **0.18** | **0.24** | **-95.21** |
| **Dist + $Depth^{30}$ + Year** | **5** | **201.18** | **0.60** | **0.19** | **-95.42** |
| **Dist + $Depth^{15}$ + Year** | **5** | **201.50** | **0.92** | **0.16** | **-95.58** |
| **Dist + $Depth^{5}$ + Year** | **5** | **201.80** | **1.23** | **0.14** | **-95.73** |

### G. Full model selection for islet physical characteristics and main prey densities, or year, on nest survival of cackling goose, glaucous gull and artificial nest

#### Cackling goose nests

Table G1. Generalized linear model selection of the effects of distance to shore (Dist) and main prey densities (lemmings (Lem) and snow goose nest densities (SnowG) on the probability of nest survival for cackling goose nesting on islets (2018, 2019, 2022 and 2023) on Bylot Island (N = 67). The exponents in the Dist variables indicate the selected decay functions. All candidate models are presented with their number of parameters (K), the change in AICc from the best-supported model (ΔAICc), Akaike weight (W), and Log-Likelihood (LogLik). Competitive models (ΔAICc < 2) are illustrated in bold.

| Models | K | AICc | ΔAICc | W | LogLik |
|---|---|---|---|---|---|
| **$Dist^{45}$ + SnowG** | **3** | **80.55** | **0.00** | **0.48** | **-37.09** |
| **SnowG** | **2** | **81.42** | **0.87** | **0.31** | **-38.62** |
| **Dist + SnowG** | **3** | **82.30** | **1.75** | **0.20** | **-37.96** |
| $Dist^{1}$ + Lem | 3 | 109.00 | 28.45 | 0.00 | -51.31 |
| Lem | 2 | 110.17 | 29.62 | 0.00 | -52.99 |
| Dist + Lem | 3 | 111.84 | 31.29 | 0.00 | -52.73 |
| Null | 1 | 123.23 | 42.67 | 0.00 | -60.58 |
| $Dist^{1}$ | 2 | 123.59 | 43.04 | 0.00 | -59.70 |
| $Dist^{45}$ | 2 | 124.52 | 43.97 | 0.00 | -60.17 |
| Dist | 2 | 125.30 | 44.75 | 0.00 | -60.56 |

Table G2. Coefficient estimates of competitive models from the generalized linear model selection of the effects of distance to shore (Dist) and main prey densities (lemmings and snow goose nests (SnowG)) on the probability of nest survival for cackling goose nesting on islets (2018, 2019, 2022 and 2023) on Bylot Island (N = 67). The exponents in the Dist variables indicate the selected decay functions. The coefficients of both Euclidean and Decay variables are presented under the same column. Coefficients' 95% confidence intervals are shown between square brackets.

| **Models** | **Int.** | **Dist** | **SnowG** |
|---|---|---|---|
| $Dist^{45}$ + SnowG | 0.53<br>[-1.29;1.58] | 10.45<br>[-3.36;48.07] | 0.05<br>[0.03;0.08] |
| SnowG | 0.96<br>[-0.37;1.94] | | 0.05<br>[0.03;0.07] |
| Dist + SnowG | 0.37<br>[-1.94;1.85] | 0.04<br>[-0.06;0.18] | 0.05<br>[0.03;0.08] |

### Glaucous gull nests

Table G3. Generalized linear model selection of the effects of distance to shore (Dist), water depth (Depth) and main prey densities (lemmings (Lem) and snow goose nests (SnowG)) on the probability of nest survival for glaucous gulls nesting islets (2018, 2019, 2022 and 2023) on Bylot Island (N = 55). The exponents in the Dist and Depth variables indicate the selected decay functions. All candidate models are presented with their number of parameters (K), the change in AICc from the best-supported model (ΔAICc), Akaike weight (W), and Log-Likelihood (LogLik). Competitive models (ΔAICc < 2) are illustrated in bold.

| Models | K | AICc | ΔAICc | W | LogLik |
|---|---|---|---|---|---|
| **Dist + SnowG** | **3** | **76.72** | **0.00** | **0.27** | **-35.13** |
| **Dist$^1$ + SnowG** | **3** | **77.17** | **0.45** | **0.21** | **-35.35** |
| **SnowG** | **2** | **78.34** | **1.62** | **0.12** | **-37.06** |
| Dist + Depth$^7$ + SnowG | 4 | 78.89 | 2.17 | 0.09 | -35.05 |
| Dist + Depth + SnowG | 4 | 79.02 | 2.30 | 0.08 | -35.11 |
| Dist$^1$ + Depth$^7$ + SnowG | 4 | 79.07 | 2.35 | 0.08 | -35.14 |
| Dist$^1$ + Depth + SnowG | 4 | 79.39 | 2.67 | 0.07 | -35.3 |
| Depth$^7$ + SnowG | 3 | 80.52 | 3.80 | 0.04 | -37.03 |
| Depth + SnowG | 3 | 80.58 | 3.86 | 0.04 | -37.05 |
| Dist$^1$ | 2 | 89.91 | 13.19 | 0.00 | -42.84 |
| Dist$^1$ + Lem | 3 | 91.06 | 14.34 | 0.00 | -42.3 |
| Dist | 2 | 91.43 | 14.71 | 0.00 | -43.6 |
| Dist$^1$ + Depth$^7$ | 3 | 91.71 | 14.99 | 0.00 | -42.62 |
| Dist + Lem | 3 | 91.93 | 15.21 | 0.00 | -42.73 |
| Dist$^1$ + Depth | 3 | 92.08 | 15.36 | 0.00 | -42.81 |
| Dist$^1$ + Depth$^7$+ Lem | 4 | 93.16 | 16.44 | 0.00 | -42.18 |
| Dist$^1$ + Depth + Lem | 4 | 93.38 | 16.66 | 0.00 | -42.29 |
| Dist + Depth$^7$ | 3 | 93.39 | 16.67 | 0.00 | -43.46 |
| Null | 1 | 93.56 | 16.84 | 0.00 | -45.74 |
| Dist + Depth | 3 | 93.65 | 16.93 | 0.00 | -43.59 |
| Lem | 2 | 94.16 | 17.44 | 0.00 | -44.97 |
| Dist + Depth$^7$ + Lem | 4 | 94.21 | 17.49 | 0.00 | -42.71 |
| Dist + Depth + Lem | 4 | 94.24 | 17.52 | 0.00 | -42.72 |
| Depth | 2 | 95.66 | 18.94 | 0.00 | -45.71 |
| Depth$^7$ | 2 | 95.68 | 18.96 | 0.00 | -45.73 |

| | | | | | |
|---|---|---|---|---|---|
| Depth + Lem | 3 | 96.28 | 19.56 | 0.00 | -44.90 |
| Depth$^{7}$ + Lem | 3 | 96.40 | 19.68 | 0.00 | -44.97 |

Table G4. Coefficient estimates of competitive models from the generalized linear model selection of the effects of distance to shore (Dist), water depth (Depth) and main prey densities (lemmings and snow goose nests (SnowG) on the probability of nest survival for glaucous gulls nesting islets (2018, 2019, 2022 and 2023) on Bylot Island (N = 55). The exponents in the Dist and Depth variables indicate the selected decay functions. The coefficients of both Euclidean and Decay variables are presented under the same column. Coefficients' 95% confidence intervals are shown between square brackets.

| **Models** | **Int.** | **Dist** | **SnowG** |
|---|---|---|---|
| Dist + SnowG | 1.07 [-0.63;2.41] | 0.07 [-0.01;0.17] | 0.03 [0.01;0.05] |
| Dist$^{1}$ + SnowG | -2.75 [-12.87;0.37] | 5.38 [1.59;16.48] | 0.02 [0.01;0.05] |
| SnowG | 2.09 [1.05;3.06] | | 0.02 [0.01;0.05] |

### Artificial nests

Table G5. Generalized linear model selection of the effects of distance to shore (Dist), water depth (Depth) and year on the probability of nest survival for artificial nests located on islets (2022-2024) on Bylot Island (N = 179). The exponents in the Dist and Depth variables indicate the selected decay functions. All candidate models are presented with their number of parameters (K), the change in AICc from the best-supported model (ΔAICc), Akaike weight (W), and Log-Likelihood (LogLik). Competitive models (ΔAICc < 2) are illustrated in bold.

| Models | K | AICc | ΔAICc | W | LogLik |
|---|---|---|---|---|---|
| **$Dist^{1}$ + Year** | **4** | **197.35** | **0.00** | **0.26** | **-94.56** |
| **$Dist^{1}$ + Depth + Year** | **5** | **198.15** | **0.80** | **0.18** | **-93.90** |
| **$Dist^{1}$ + $Depth^{75}$ + Year** | **5** | **198.15** | **0.80** | **0.17** | **-93.90** |
| Dist + Year | 4 | 200.04 | 2.69 | 0.07 | -95.91 |
| Dist + $Depth^{75}$ + Year | 5 | 200.58 | 3.23 | 0.05 | -95.12 |
| Dist + Depth + Year | 5 | 200.69 | 3.34 | 0.05 | -95.17 |
| $Dist^{1}$ + $Depth^{75}$ + Year + $Depth^{75}$:Year | 7 | 200.79 | 3.44 | 0.05 | -93.07 |
| $Dist^{1}$ + Depth + Year + Depth:Year | 7 | 200.83 | 3.48 | 0.05 | -93.09 |
| $Dist^{1}$ + $Depth^{75}$ + Year + $Dist^{1}$:Year | 7 | 201.68 | 4.33 | 0.03 | -93.51 |
| $Dist^{1}$ + Depth + Year + $Dist^{1}$:Year | 7 | 201.69 | 4.34 | 0.03 | -93.52 |
| Dist + $Depth^{75}$ + Year + $Depth^{75}$:Year | 7 | 203.29 | 5.94 | 0.01 | -94.32 |
| Dist + $Depth^{75}$ + Year + Dist:Year | 7 | 203.44 | 6.09 | 0.01 | -94.39 |
| Dist + Depth + Year + Depth:Year | 7 | 203.50 | 6.14 | 0.01 | -94.42 |
| Dist + Depth + Year + Dist:Year | 7 | 203.59 | 6.24 | 0.01 | -94.47 |
| Year | 3 | 204.23 | 6.88 | 0.01 | -99.05 |
| $Depth^{75}$ + Year | 4 | 205.56 | 8.21 | 0.00 | -98.67 |
| Depth + Year | 4 | 205.81 | 8.46 | 0.00 | -98.79 |
| Null | 1 | 208.13 | 10.78 | 0.00 | -103.06 |

Table G6. Coefficient estimates of competitive models from the generalized linear model selection of the effects of distance to shore (Dist), water depth (Depth) and year on the probability of nest survival for artificial nests located on islets (2022-2024) on Bylot Island (N = 179). The exponents in the Dist and Depth variables indicate the selected decay functions. The coefficients of both Euclidean and Decay variables are presented under the same column. Coefficients' 95% confidence intervals are presented between square brackets.

| **Models** | **Int.** | **Dist** | **Depth** | **$Year_{2023}$** | **$Year_{2024}$** |
|---|---|---|---|---|---|
| $Dist^{1}$ + Year | -7.23 [-13.12;-3.86] | 6.29 [2.38;12.84] | | 1.24 [0.44;2.41] | 1.26 [0.27;2.48] |
| $Dist^{1}$ + Depth + Year | -7.09 [-13.09;-3.71] | 6.73 [2.77;13.51] | -0.02 [-0.05;0.01] | 1.18 [0.33;2.32] | 1.30 [0.31;2.54] |
| $Dist^{1}$ + $Depth^{75}$ + Year | -7.24 [-13.24;-3.94] | 6.60 [2.71;13.27] | -2.89 [-9.17;1.84] | 1.19 [0.34;2.29] | 1.30 [0.33;2.49] |
